# The ciliary protein C2cd3 is required for mandibular musculoskeletal tissue patterning

**DOI:** 10.1101/2024.02.23.581730

**Authors:** Evan C. Brooks, Simon J.Y. Han, Christian Louis Bonatto Paese, Amya A. Lewis, Megan Aarnio-Peterson, Samantha A. Brugmann

## Abstract

The mandible is composed of several musculoskeletal tissues including bone, cartilage, and tendon that require precise patterning to ensure structural and functional integrity. Interestingly, most of these tissues are derived from one multipotent cell population called cranial neural crest cells (CNCCs). How CNCCs are properly instructed to differentiate into various tissue types remains nebulous. To better understand the mechanisms necessary for the patterning of mandibular musculoskeletal tissues we utilized the avian mutant *talpid^2^* (*ta^2^*) which presents with several malformations of the facial skeleton including dysplastic tendons, mispatterned musculature, and bilateral ectopic cartilaginous processes extending off Meckel’s cartilage. We found an ectopic epithelial BMP signaling domain in the *ta^2^* mandibular prominence (MNP) that correlated with the subsequent expansion of *SOX9*+ cartilage precursors. These findings were validated with conditional murine models suggesting an evolutionarily conserved mechanism for CNCC-derived musculoskeletal patterning. Collectively, these data support a model in which cilia are required to define epithelial signal centers essential for proper musculoskeletal patterning of CNCC-derived mesenchyme.

## Introduction

The mandible is an anatomically complex structure composed of numerous tissue types, including cartilage, bone, muscle, tendons, ligaments, nerves, and vasculature, that allows for essential behaviors such as communication and mastication (Klingenberg and Navarro, 2012). Each tissue component of the mandible must be temporally and spatially patterned, and differentiated in concert, to ensure structural and functional integrity. Disruptions and/or variations in mandibular development can result in several musculoskeletal conditions and negatively impact patients’ ability to speak, breath and chew.

Development of the mandible begins when multipotent cranial neural crest cells (CNCCs) are generated in the dorsal neural tube. CNCCs delaminate from the neural tube and subsequently migrate into the mandibular prominence (MNP) (Couly et al., 1998; Köntges and Lumsden, 1996). As CNCCs migrate into the MNP, they are exposed to numerous signals from the surrounding epithelium including *Sonic hedgehog* (*Shh*) from the pharyngeal endoderm and oral ectoderm, *Bone morphogenetic protein 4* (*Bmp4*) from the distal aboral ectoderm, and *Fibroblast growth factor 8* (*Fgf8*) from the proximal aboral ectoderm (Bitgood and McMahon, 1995; Tucker et al., 1999; Vainio et al., 1993; Xu et al., 2019). These instructive signals regulate the expression of numerous downstream transcription factors that ultimately control the differentiation of CNCCs into numerous musculoskeletal tissues (e.g., bone, cartilage, tendon). Shh is necessary for the survival of post-migratory CNCCs and the subsequent differentiation of chondrocytes (Billmyre and Klingensmith, 2015). CNCCs that receive a Bmp signal express *SRY-box transcription factor 9* (*Sox9*) and subsequently differentiate into chondrocytes to form Meckel’s cartilage, the structural scaffold for the developing mandible (Bi et al., 1999; Hu et al., 2008; Svandova et al., 2020). CNCCs that receive an Fgf signal will express the transcription factor *Scleraxis* (*Scx*) and will differentiate into tenocytes and ligamentocytes (Bobzin et al., 2021; Edom-Vovard et al., 2002; Havis et al., 2016). Despite having knowledge of the initial signals that prompt CNCC differentiation into numerous musculoskeletal lineages, a complete understanding of how these signals integrate to generate a functional mandible, and how disruptions in these signaling pathways cause syndromes affecting mandibular development, is incomplete.

Mandibular development is frequently disrupted in a class of diseases called ciliopathies, which arise from disruptions in the structure and function of the primary cilium - a microtubule-based cellular extension that functions as a centralized hub for molecular signal transduction (Elliott and Brugmann, 2019; Goetz and Anderson, 2010). Ciliopathies affect as many as 1 in 800 human patients and are characterized by numerous pleiotropic phenotypes including skeletal dysplasias and craniofacial anomalies (Baker and Beales, 2009; Reiter and Leroux, 2017). Approximately 30% of all ciliopathies can be classified by their craniofacial phenotypes, including cleft lip/palate, craniosynostosis, and micrognathia (Schock and Brugmann, 2017). Human patients and animal models that present with craniofacial ciliopathies have an array of mandibular phenotypes such as micrognathia, dysmorphic skeletal elements, and reduced bone density (Adel Al-Lami et al., 2016; Bonatto Paese et al., 2021a; Cela et al., 2018; Ferrante et al., 2009; Kantaputra et al., 2023; Kitamura et al., 2020; Kolpakova-Hart et al., 2007; Zhang et al., 2011). While significant progress has been made in understanding the molecular etiologies of craniofacial ciliopathies, comprehensive knowledge of how individual ciliary proteins regulate the spatial patterns of early molecular signals that contribute to musculoskeletal tissue patterning is currently lacking.

Oral-facial-digital syndrome (OFD) is an umbrella term for at least 18 distinctive ciliopathic disorders characterized by anomalies in the oral cavity, facial structures, and digits (Franco and Thauvin-Robinet, 2016). Several types of OFD present with mandibular phenotypes including micrognathia, retrognathia, hypo-or aglossia, hamartomas, and ankyloglossia. OFD14 is caused by mutations in C2 domain containing 3 centriole elongation regulator (C2CD3), which is localized to the distal centrioles and acts as a critical elongation factor during ciliogenesis (Boczek et al., 2018; Cortés et al., 2016; Hoover et al., 2008; Thauvin-Robinet et al., 2014; Ye et al., 2014). To study the molecular etiologies of phenotypes associated with human OFD14, we utilized the *talpid^2^* (*ta^2^*). The *ta^2^* is a naturally occurring avian mutant that is characterized by its numerous craniofacial phenotypes as well as its polydactylous limbs (Abbott et al., 1960, 1959; Bonatto Paese et al., 2022, 2021a; Brugmann et al., 2010; Caruccio et al., 1999; Chang et al., 2014; Schock et al., 2015). We previously reported that the *ta^2^* phenotype was the result of a 19 bp deletion in *C2CD3* (Chang et al., 2014). Interestingly, this *C2CD3* variant faithfully phenocopies human OFD14 (Bonatto Paese et al., 2021a; Chang et al., 2014; Harris et al., 2006; Schock et al., 2015), presenting with strikingly similar craniofacial, neural, and limb phenotypes, making the *ta^2^* an excellent model for understanding the molecular etiology for several craniofacial and musculoskeletal phenotypes.

Herein, we characterized previously unreported ectopic cartilaginous processes extending off Meckel’s cartilage in *ta^2^* embryos. The generation of these structures correlated with an ectopic, epithelial BMP signaling center and increased expression of BMP target genes. Together, these results suggest a novel cellular and molecular mechanism for musculoskeletal craniofacial phenotypes present in OFD14.

## Materials and Methods

### Avian embryo collection, processing, and genotyping

Fertilized control^+/+^ and *ta^2^* eggs were supplied by the University of California, Davis Avian Facility. Eggs were incubated at 38.8°C for 2 – 13 days in a rocking incubator with humidity control. Embryos were staged according to the Hamburger-Hamilton (HH) staging system (Hamburger and Hamilton, 1951), harvested in cold diethyl pyrocarbonate-treated phosphate-buffered solution (DEPC PBS) and fixed in either 10% neutral buffered formalin (NBF) overnight at room temperature or 4% paraformaldehyde (PFA) in DEPC PBS overnight at 4°C. Embryos were genotyped as previously described (Chang et al., 2014).

### Mouse strains

All mouse strains used in this study have been previously described: *AP2-Cre* (Macatee et al., 2003), *C2cd3^ex4-5flox^* (Chang et al., 2021), *Crect* (Reid et al., 2011), *Scx-Cre* (Blitz et al., 2009), and *Wnt1-Cre2* (Lewis et al., 2013). Both male and female mice were used. A maximum of 4 adult mice were housed per cage and breeding cages housed one male paired with up to two females. All mouse usage was approved by the Institutional Animal Care and Use Committee (IACUC) and monitored daily by the Division of Veterinary Services at Cincinnati Children’s Hospital Medical Center.

### Mouse embryo collection and genotyping

Timed matings were performed, with noon of the day a vaginal plug was discovered designated as E0.5. Embryos were harvested via Caesarian section after isoflurane-induced asphyxiation and cervical dislocation, and were dissected in cold DEPC PBS. Samples were fixed in 10% NBF overnight at room temperature or 4% PFA in DEPC PBS overnight at 4°C. For all experiments, *Cre*-negative littermates were used as controls. Genotyping was performed using the primers in Table 1.

**Table 1.**
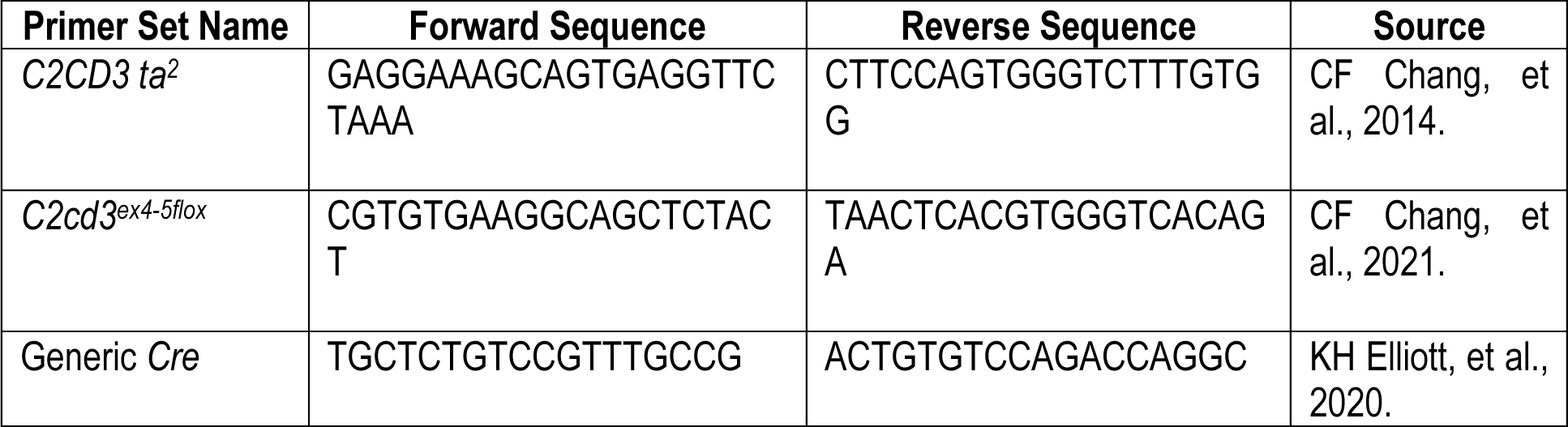
Genotyping Primers.

### Wholemount skeletal staining and morphometrics

HH30-32 avian embryos were stained with .05% Alcian blue solution to detect cartilage and cleared in 1:2 benzyl alcohol:benzyl benzoate solution as previously described (Nagy et al., 2009). HH34-39 avian embryos and E15.5-E17.5 mouse embryos were first immersed in hot water to facilitate the manual removal of skin and soft tissues and were stained with .015% Alcian blue solution to detect cartilage. E17.5 mouse embryos were further stained for bone with 0.005% Alizarin red in 1% KOH solution. Samples were then cleared in graded 1% KOH/glycerol solutions and imaged in 50% glycerol as previously described (Elliott et al., 2020). All samples were imaged using a Leica M165 FC stereo microscope system. Area measurements were performed by tracing the outline of Meckel’s cartilage on HH30 and HH36 avian embryos via the freehand tool in FIJI (Schindelin et al., 2012).

### Wholemount hybridization chain reaction

The Hybridization Chain Reaction (HCR) assay from Molecular Instruments, Inc. was carried out as previously described (Choi et al., 2018, 2016). HCR on HH14-20 avian mandibular prominences and e10.5 mouse mandibular prominences was performed using the manufacturer’s instructions for chicken and mouse embryos, respectively. For HCR assays performed on HH36 avian mandibles, several modifications were made. Samples were quickly blanched in a 55°C water bath shortly after fixation to facilitate the dissection of excess keratinized epithelium to enhance reagent penetration, proteinase K solution concentration was increased to 50 μg/mL, and all PBS with 0.1% Tween and 5× SSC with 0.1% Tween (SSCT) washes were increased to 10 min. All probes used in this study were designed by Molecular Instruments and can be found in Table 2. All samples were counterstained with 1:10000 dilution of Hoescht 33342 solution (Invitrogen H3570) in 5× SSCT for 5 min at room temperature then washed five times in 5× SSCT. Samples were then cleared in multiple changes of refractive index matching solution (Muntifering et al., 2018) at room temperature until samples were transparent. Samples were imaged on either a Nikon FN1 upright confocal microscope system or a Nikon A1 inverted confocal microscope system. Images were denoised in Nikon NIS-Elements software and then loaded into Imaris 10.0.0 software for analysis and visualization. Three-dimensional renderings of avian and murine *SHH*, *BMP4,* and *FGF8* expression data were generated using Imaris 10.0 software via the Surfaces Tool.

**Table 2.** HCR Probes.

| Probe Name | Species | Channel | GenBank Identifier | Lot Number |
| --- | --- | --- | --- | --- |
| *BMP4* | *Gallus gallus* (chicken) | B3 | NM_205237.4 | PRO952 |
| *COL2A1* | *Gallus gallus* (chicken) | B5 | NM_204426.2 | RTD418 |
| *FGF8* | *Gallus gallus* (chicken) | B1 | NM_001012767.2 | PRK963 |
| *MYOG* | *Gallus gallus* (chicken) | B4 | NM_204184.2 | RTG675 |
| *SCX* | *Gallus gallus* (chicken) | B1 | NM_204253.2 | PRO152 |
| *SHH* | *Gallus gallus* (chicken) | B2 | NM_204821.1 | RTE050 |
| *Bmp4* | *Mus musculus* (mouse) | B3 | NM_007554.3 | RTC282 |
| *Fgf8* | *Mus musculus* (mouse) | B2 | NM_010205 | RTC512 |
| *Shh* | *Mus musculus* (mouse) | B1 | NM_009170.3 | RTC686 |

### Paraffin embedding and sectioning

Samples were dehydrated in a graded ethanol series, washed in xylene, embedded in paraffin, and sectioned at 6 – 8 μm thickness using a Sakura Accu-Cut SRM 200 microtome. Sections were dried overnight at room temperature.

### RNAscope in situ hybridization

The RNAscope^®^ assay was carried out as previously described (Bonatto Paese et al., 2021a; Wang et al., 2012). Briefly, transcripts were detected using the RNAscope Multiplex Fluorescent V2 kit per manufacturer’s instructions. All probes to detect transcripts used in this study were designed by Advanced Cell Diagnostics and can be found in Table 3. Signals for the transcripts were detected using fluorescein (Akoya Biosciences NEL741001KT) and Cyanine 3 (Akoya Biosciences NEL744001KT) diluted 1:500 in RNAscope Multiplex TSA Buffer. All samples were counterstained with RNAscope DAPI (4’,6-diamidino-2-phenylindole) solution and imaged on a Leica DM5000B upright microscope system.

**Table 3.**
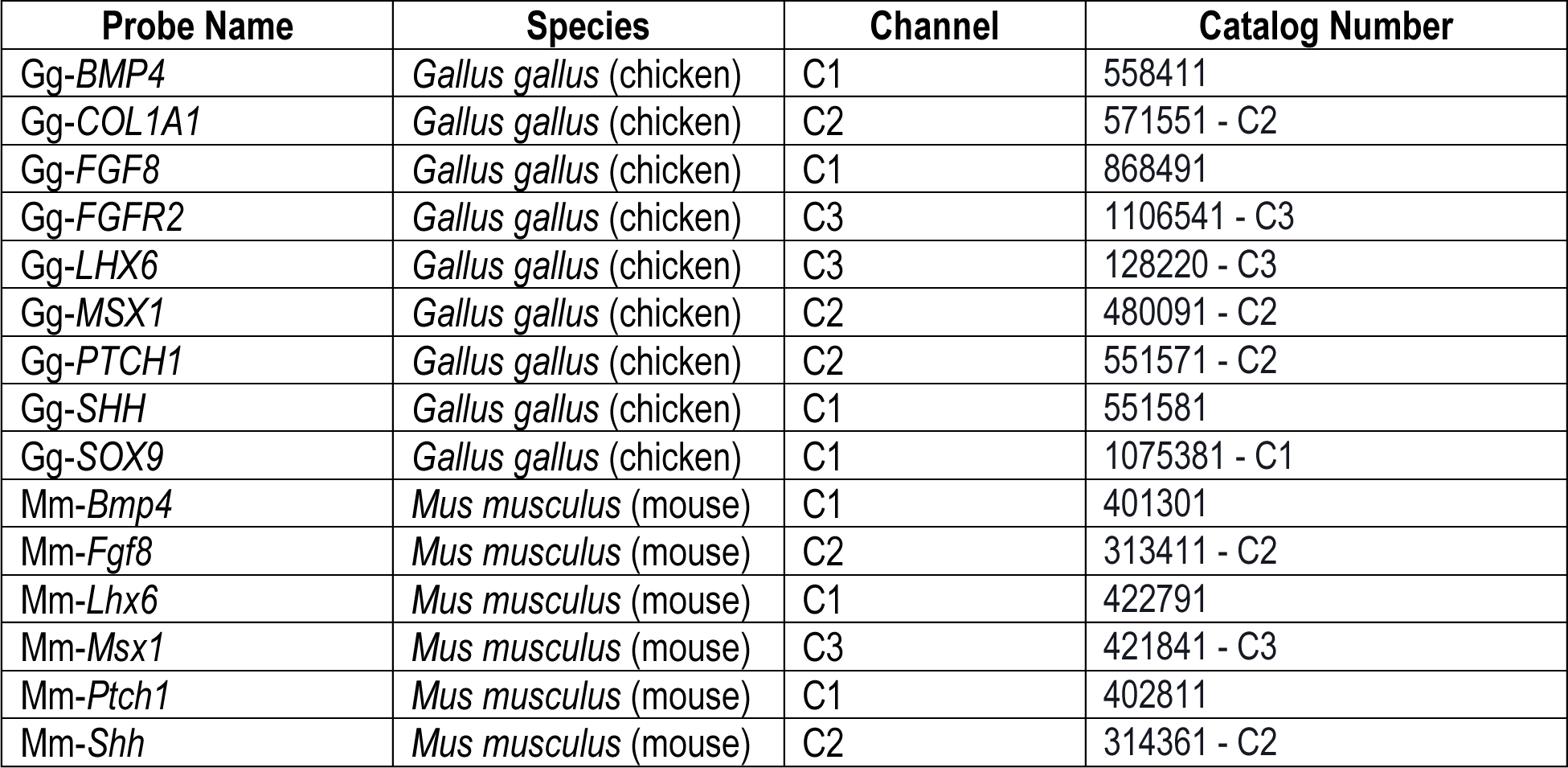
RNAscope Probes.

| Probe Name | Species | Channel | Catalog Number |
| --- | --- | --- | --- |
| Gg-BMP4 | *Gallus gallus* (chicken) | C1 | 558411 |
| Gg-COL1A1 | *Gallus gallus* (chicken) | C2 | 571551 - C2 |
| Gg-FGF8 | *Gallus gallus* (chicken) | C1 | 868491 |
| Gg-FGFR2 | *Gallus gallus* (chicken) | C3 | 1106541 - C3 |
| Gg-LHX6 | *Gallus gallus* (chicken) | C3 | 128220 - C3 |
| Gg-MSX1 | *Gallus gallus* (chicken) | C2 | 480091 - C2 |
| Gg-PTCH1 | *Gallus gallus* (chicken) | C2 | 551571 - C2 |
| Gg-SHH | *Gallus gallus* (chicken) | C1 | 551581 |
| Gg-SOX9 | *Gallus gallus* (chicken) | C1 | 1075381 - C1 |
| Mm-Bmp4 | *Mus musculus* (mouse) | C1 | 401301 |
| Mm-Fgf8 | *Mus musculus* (mouse) | C2 | 313411 - C2 |
| Mm-Lhx6 | *Mus musculus* (mouse) | C1 | 422791 |
| Mm-Msx1 | *Mus musculus* (mouse) | C3 | 421841 - C3 |
| Mm-Ptch1 | *Mus musculus* (mouse) | C1 | 402811 |
| Mm-Shh | *Mus musculus* (mouse) | C2 | 314361 - C2 |

### Avian embryo paralysis

Decamethonium bromide (DBr) (Sigma-Aldrich D1260) was diluted to 10 mg/ml in Hank’s Buffered Sterile Saline (HBSS) and filter sterilized using a 0.22 μm filter. HH29 embryos were treated with 0.5 mL DBr via injection into the albumin with a syringe through a pinhole made in the eggshell as previously described (Hall, 1975; Woronowicz et al., 2018). Embryos were assayed for paralysis via hindlimb extension before collecting for Alcian blue staining.

### Single-cell RNA-sequencing

Three mandibular prominences per group from HH26 control and *talpid^2^* embryos were quickly dissected in ice-cold DEPC PBS. Any regions rich in blood were manually removed. The MNPs were then minced to a fine paste. Cells were dissociated into a single-cell suspension and sequenced using NovaSeq 6000 and the S2 flow cell. 12.5 mg of tissue was placed in a sterile 1.5 mL tube containing 0.5 mL protease solution containing 125 U/mL DNase and 3 mg/mL *Bacillus Licheniformis*. The samples were incubated on ice for a total of 10 min, with gentile trituration using a wide boar pipette tip every minute after the first two. Protease was inactivated using ice-cold DEPC PBS containing 0.02% BSA and filtered using 30 µM filter. The cells were pelleted by centrifugation at 200G for 4 min at room temperature and resuspended in 50 μL Dulbecco’s Modified Eagle Medium with 10% Fetal Bovine Serum. Cell number and viability were assessed using a hemocytometer and trypan blue staining. 9,600 cells were loaded onto a well on a 10x Chromium Single-Cell instrument (10X Genomics) to target sequencing of 6,000 cells. Barcoding, cDNA amplification, and library construction were performed using the Chromium Single-Cell 3’ Library and Gel Bead Kit v3. Post cDNA amplification and cleanup was performed using SPRI select reagent (Beckman Coulter B23318). Post cDNA amplification and post-library construction quality control was performed using the Agilent Bioanalyzer High Sensitivity kit (Agilent 5067–4626). Libraries were sequenced using a NovaSeq 6000 and the S2 flow cell. Sequencing parameters used were: Read 1, 28 cycles; Index i7, eight cycles; Read 2, 91 cycles, producing about 300 million reads. Sequenced reads were mapped to the Ensembl build of the chicken genome GRCg6a using CellRanger (http://10xgenomics.com) to obtain a gene-cell data matrix.

### Analysis of single-cell RNA-sequencing data

Analysis of the data was conducted using the Seurat v4.4.0 package (Hao et al., 2021). Control and *ta^2^* cells were filtered for quality by unique feature counts (less than 200 or greater than 4000) and mitochondrial gene expression (greater than 15%) to remove any low-quality cells. Filtered cells were then normalized and cell cycle genes were removed. “RunPCA,” “RunUMAP,” and “FindNeighbors” were then performed using 20 dimensions along with “FindClusters” with a resolution of 1 to generate the working R objects. The control and *ta^2^* datasets were then integrated by “FindIntegrationAnchors” and “IntegrateData” and the same filtering criteria were applied to generate the working integrated R object.

### CellChat analysis

The standard CellChat workflow was used to assay signaling interactions as previously described (Jin et al., 2021). In brief, the individual Seurat objects were used as inputs for CellChat. These objects were processed by the standard parameters using the human CellChat database as the reference. Once the top pathways list was generated, the functions “plotGeneExpression”, “netAnalysis_computeCentrality”, “netAnalysis_signalingRole_network”, and “netVisual_aggregate” were used to generate violin plots, heatmaps, and circle plots of the pathways of interest.

### Micro-computed tomography of murine embryo heads

Heads from E17.5 *C2cd3^ex4-5fl/fl^* and *C2cd3^ex4-5fl/fl^; AP2-Cre* embryos were harvested in cold DEPC PBS and fixed overnight in 4% PFA at 4°C. Microcomputed tomography (microCT) was performed using a Siemens Inveon PET/SPECT/CT scanner in the Preclincal Imaging Core of the University of Cincinnati Vontz Center for Molecular Studies. The cone-beam CT parameters were as follows: 360° rotation, 1080 projections, 1300 ms exposure time, 1500 ms settle time, 80 kVp voltage, 500 μA current, and effective pixel size 17.67 μm. Acquisitions were reconstructed using a Feldkamp algorithm with slight noise reduction, 3D matrix size 1024×1024×1536, using manufacturer-provided software. Protocol-specific Hounsfield Unit (HU) calibration factor was applied. DICOM files were loaded into 3D Slicer 5.6.0 software (Fedorov et al., 2012). Three-dimensional renderings of the head were constructed, surface smoothing was executed, and the mandible was segmented via the Segment Editor function. Mandibular length measurements were taken from the mandibular symphysis to the condylar process as previously described (Roberts et al., 2019) on both sides of the mandible and averaged. Ectopic mandibular skeletal elements were manually pseudocolored in Adobe Illustrator.

### Statistical methods

Unpaired two-tailed, Student’s *t*-test were used in all statistical analyses performed at the 5% significance level. P-values less than .05 (*p* < .05) were considered statistically significant.

## Results

### ta^2^ embryos presented with ectopic, bilateral cartilaginous processes extending off Meckel’s cartilage

Previous reports from our lab and others have documented the craniofacial and skeletal phenotypes present in *ta^2^* embryos (Abbott et al., 1960, 1959; Bonatto Paese et al., 2022, 2021b, 2021a; Brooks et al., 2021; Brugmann et al., 2010; Caruccio et al., 1999; Chang et al., 2014; Dvorak and Fallon, 1991; Harris et al., 2006; Schneider et al., 1999; Schock et al., 2015). To better understand the molecular etiology of these phenotypes, we performed a temporal analysis of mandibular development, initially focused on Meckel’s cartilage. Alcian blue staining and measurements of HH30 control and *ta^2^* Meckel’s cartilages revealed no discernable phenotypic or size difference (Fig. 1A-B, Fig. S1A). Between HH32-34, however, ectopic, bilateral cartilaginous processes were consistently observed extending from the more distal aspect of Meckel’s cartilage (Fig. 1C-F). By HH36, these ectopic processes were prominent (Fig, 1G-H) and contributed to an increase in the total area of Meckel’s cartilage (Fig. S1B). The ectopic processes were consistently observed with a conserved shape and location until embryonic lethality at HH39 (Fig. 1I-J) (Abbott et al., 1960, 1959).

**Figure 1.**
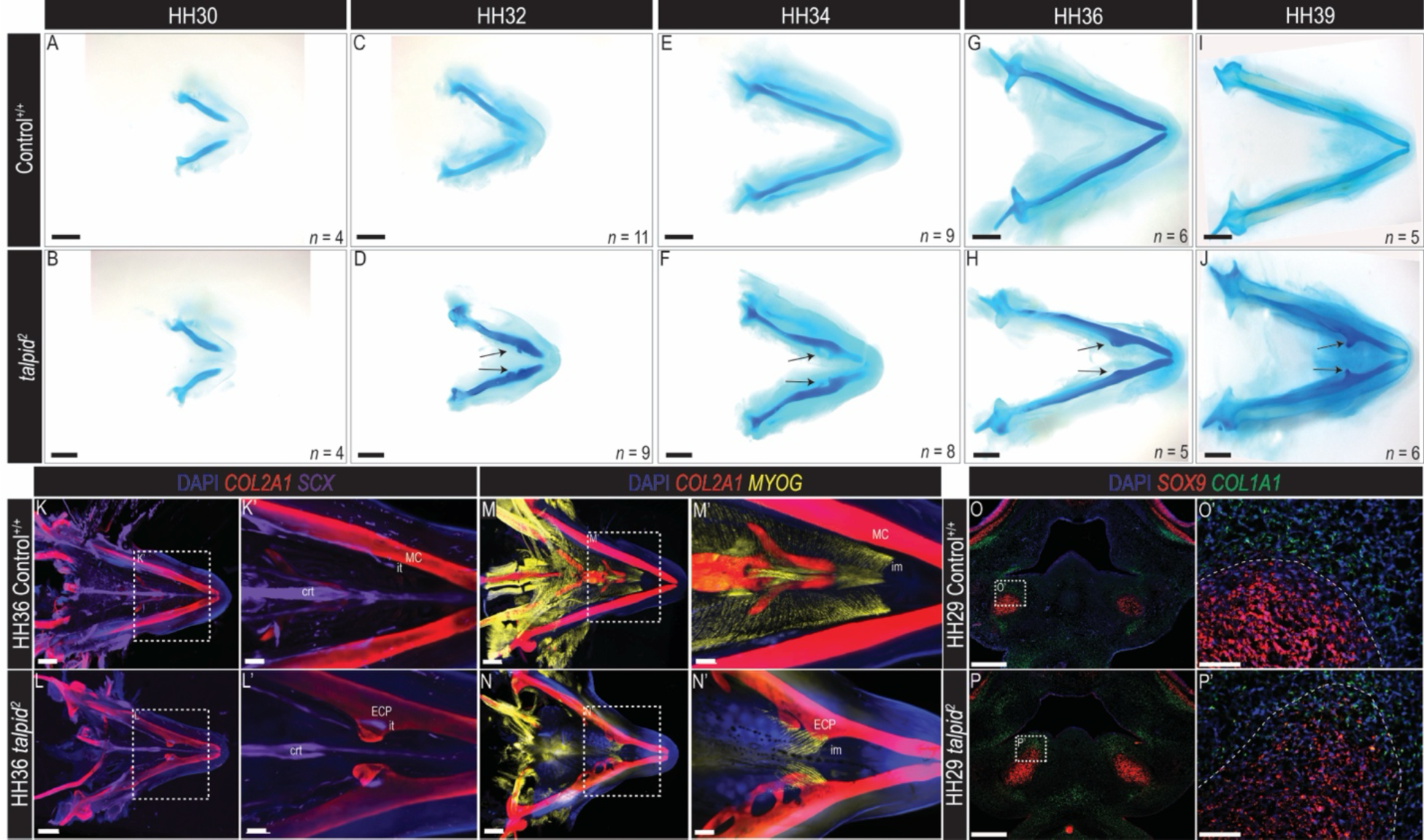
*ta^2^* mandibles present with numerous musculoskeletal anomalies reminiscent of enthesis formation. (A-J) Dorsal views of Alcian blue-stained HH30 (A-B), HH32 (C-D), HH34 (E-F), HH36 (G-H), and HH39 (I-J) control^+/+^ (A, C, E, G, I) and *ta^2^* (B, D, F, H, J) Meckel’s cartilages. (K-N’) Ventral views of wholemount hybridization chain reaction (HCR) assays for *SOX9* and *SCX* to detect cartilage (red) and tendon (purple) (K-L’) or *COL2A1* and *MYOG* to detect cartilage (red) and muscle (yellow) (M-N’) in HH36 control^+/+^ (K-K’, M-M’) and *ta^2^* (L-L’, N-N’) mandibles (*n* = 3 per group). (O-P’) Section RNAscope *in situ* hybridization assays for *SOX9* and *COL1A1* to detect presumptive cartilage (red) and connective tissues (green) in frontal sections of HH29 control^+/+^ (O-O’) and *ta^2^* (P-P’) mandibular prominences (*n* = 3 per group). Dotted lines in O’ and P’ denote the outline of Meckel’s cartilage as denoted by *SOX9* expression. Arrowheads in D, F, H, and J denote *ta^2^* ectopic cartilaginous process. Dotted boxes in K-P denote areas of higher magnification. crt: central raphe tendon, ECP: ectopic cartilaginous process, im: intermandibular muscle, it: intermandibular tendon, MC: Meckel’s cartilage. Scale bars: 1mm (A-K, L, M, N); 300μm (K’, L’, M’, N’); 500μm (O, P); 50μm (O’, P’).

The presence of these ectopic cartilaginous processes was surprising, as the *ta^2^* has been studied for nearly 7 decades without mention of such structures. As such, we searched the literature to see if similar structures had previously been reported. Interestingly, *Runx2*-deficient mice were reported to have two ectopic cartilaginous processes on Meckel’s cartilage which attached to the digastric and mylohyoid muscles (Shibata et al., 2004). These data, coupled with our previous findings that mandibular skeletal elements were hypoplastic in the *ta^2^* embryos (Bonatto Paese et al., 2021a), prompted the examination of other musculoskeletal tissues. Using wholemount hybridization chain reaction (HCR; Choi et al., 2018, 2016) we examined the organization of tendons and muscle relative to cartilage in HH36 embryos via the expression of *SCLERAXIS* (*SCX*; Cserjesi et al., 1995; Schweitzer et al., 2001), *MYOGENIN* (*MYOG*; Wright et al., 1989) and *COLLAGEN TYPE II ALPHA 1 CHAIN (COL2A1*; Kosher et al., 1986), respectively. In control embryos, the intermandibular tendons ran parallel to, and were immediately adjacent to Meckel’s cartilage (Fig. 1K-K’); however, in *ta^2^* mandibles, they appeared to insert directly into the ectopic cartilaginous processes (Fig. 1L-L’). Similarly, the intermandibular muscle, which spans the entire length of Meckel’s cartilage (Fig. 1M-M’; Robson, 1993), was disorganized, hypoplastic and also appeared to insert directly into the ectopic cartilaginous processes (Fig. 1N-N’). Thus, in addition to the presence of ectopic cartilaginous processes, the *ta^2^* mandible presented with a musculoskeletal disorganization similar to other mutations that impact skeletal development of CNCCs.

The appearance and organization of ectopic cartilaginous processes and accompanying ectopic musculoskeletal attachments in the *ta^2^* mandible phenotypically resembled a secondary cartilage, or a cartilage that is independent from the primary cartilaginous skeleton and arises on existing dermal bones at sites of articulations and insertions (Hall, 2015; Murray, 1963). Secondary cartilages can be induced through mechanical forces and express both *FIBROBLAST GROWTH FACTOR RECEPTOR 2* (*FGFR2*) and *BMP4* (de Beer and Barrington, 1934; Solem et al., 2011). Given the observation that the intermandibular muscle appeared to be inserted into the ectopic cartilaginous processes, we hypothesized that they could be secondary cartilages caused by muscle contraction. To test this hypothesis, *ta^2^* embryos were paralyzed via decamethonium bromide (DBr) treatment and characterized molecularly. Neither the presentation nor size of the ectopic cartilaginous processes was alleviated via paralysis (Fig. S2A-D). Furthermore, the processes did not express *BMP4* or *FGFR2* (Fig. S2E-F’). Thus, these results suggested that ectopic cartilaginous processes observed in *ta^2^* embryos were not secondary cartilages.

Given that our molecular analysis did not support the identity of the ectopic cartilaginous processes as secondary cartilages and that the intermandibular muscle was attached to these structures, we next tested the hypothesis that they were precursors of entheses, or tendon-bone interfaces. In the stages preceding enthesis specification and mineralization, the musculature attaches to the cartilaginous template of the developing skeleton (Schweitzer et al., 2010; Sunadome et al., 2023). Molecularly, an enthesis develops from a cell population that co-expresses factors associated with cartilage (*SOX9*) and tendon (*SCX, COL1A1*) development (Blitz et al., 2013; Bobzin et al., 2021; Sugimoto et al., 2013). Utilizing RNAscope for *SOX9* and *COL1A1* in HH29 control mandibular prominences (MNPs), we found distinct areas of either *SOX9*+ or *COL1A1*+ cells that did not intermix (Fig. 1O-O’). Conversely, in *ta^2^* embryos, the domain of *SOX9*+ cells expanded into the *COL1A1*+ area resulting in several *SOX9*+/*COL1A1*+ cells (Figure 1P-P’). Thus, taken together, these analyses suggested that the ectopic cartilaginous processes present on Meckel’s cartilage molecularly and anatomically resembled an enthesis during mandibular musculoskeletal development. Considering these findings, we next sought to understand the molecular etiology of these ectopic processes in the *ta^2^*.

### Ectopic cartilage formation in ta^2^ embryos correlated with ectopic epithelial expression of BMP4

The generation of the enthesis-like structures in the *ta^2^* mandible prompted us to investigate the mechanisms that induce CNCCs to differentiate into various mandibular musculoskeletal tissues. As CNCCs migrate into the MNP, they experience signals from compartmentalized epithelial domains including *BMP4* in the aboral distal ectoderm, *FGF8* in the aboral proximal ectoderm, and *SHH* in the pharyngeal endoderm and oral ectoderm (Jeong et al., 2004; Shigetani et al., 2000; Tucker et al., 1999). These factors antagonize each other during early mandibular development to ensure proper mesenchymal gene expression and subsequent musculoskeletal tissue differentiation (Haworth et al., 2007, 2004; Neubüser et al., 1997; Tucker et al., 1998; Xu et al., 2019). Previous studies demonstrated that the combinatorial action of *FGF8* and *SHH* is required to drive cartilaginous outgrowth in the developing craniofacial complex (Abzhanov and Tabin, 2004; Hu et al., 2003) while other studies indicated that exogenous *BMP4* expression can lead to ectopic craniofacial chondrogenesis (Barlow and Francis-West, 1997; Hu et al., 2008; Nonaka et al., 1999; Semba et al., 2000). With the knowledge that primary cilia regulate signal transduction and the *ta^2^* embryo has dysregulated Hedgehog signaling in the developing craniofacial complex (Brooks et al., 2021; Brugmann et al., 2010; Chang et al., 2014), we sought to examine the relative expression patterns of *BMP4*, *FGF8* and *SHH* in control and *ta^2^* MNPs.

To uncover the potential molecular basis for ectopic cartilage in *ta^2^* embryos, we began our analysis at HH20, a developmental stage that precedes the onset of gross *ta^2^* craniofacial anomalies. In dorsal views of HH20 control MNPs, *SHH* and *BMP4* were expressed in complementary domains across the oral-aboral axis (Xu et al., 2019), whereas *FGF8* was expressed in proximal domains (Figure 2A). While *FGF8* was maintained in the proximal domain of the *ta^2^* MNP, the *SHH* domain was diminished and the *BMP4* domain was expanded (Figure 2B). Sagittal views confirmed that *BMP4* expression was expanded into the oropharyngeal domain of HH20 *ta^2^* MNPs while the *SHH* domain was diminished when compared to stage-matched controls (Figure 2C-D). Quantification of the expression domain surface area confirmed a significant decrease in the *SHH* domain and a significant increase in the *BMP4* domain with no change to the *FGF8* domain (Figure 2E). Interestingly, the *BMP4* domain was significantly increased and was expanded into the oropharyngeal aspect of the *ta^2^* MNP by HH14 (Fig. S3), a stage at which the MNP is being patterned (Abramyan and Richman, 2018; Hamburger and Hamilton, 1951).

**Figure 2.**
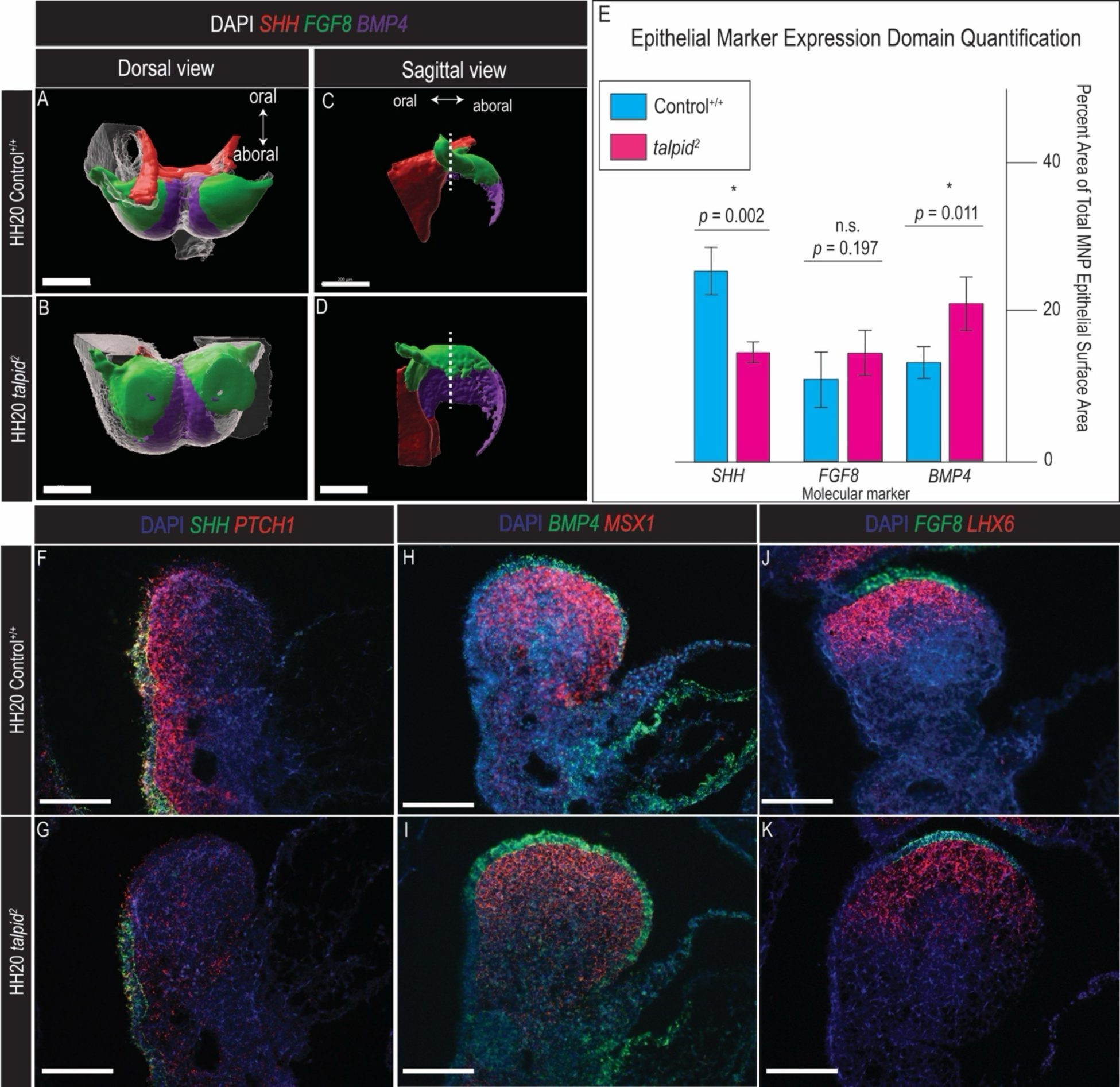
Expression of BMP signaling factors is expanded in *ta^2^* MNPs. (A-D’) Three-dimensional renderings of wholemount HCR expression data for *SHH* (red), *FGF8* (green), and *BMP4* (purple) in HH20 control^+/+^ (A, C, *n* = 4) and *ta^2^* (B, D, *n* = 4) MNPs. (E) Quantitation of three-dimensional surface renderings of *SHH*, *FGF8*, and *BMP4* in HH20 control^+/+^ and *ta^2^* MNPs as a percentage of total mandibular surface area. (F-K) RNAscope *in situ* hybridization for *SHH* and *PTCH1* (F-G), *BMP4* and *MSX1* (H-I), and *FGF8* and *LHX6* (J-K) on HH20 control^+/+^ (F, H, J) and *ta^2^* (G, I, K) sagittal MNP sections. *n* = 3 for all RNAscope experiments. The dotted lines in C and D denote the midpoint of the oral-aboral axis of the MNP. Scale bars: 200μm (A-D), 100μm (F-K). *p* > 0.05, n.s. signifies no significance.

With the observation that the spatial organization of epithelial *BMP4* and *SHH* expression was perturbed in the *ta^2^*, we next examined the expression of downstream targets in the underlying mesenchyme utilizing RNAscope *in situ* hybridization in HH20 MNPs. In HH20 control MNPs, *SHH* and its downstream transcriptional target *PATCHED1* (*PTCH1*; Goodrich et al., 1996; Marigo et al., 1996) were expressed in the oropharyngeal endoderm and adjacent mesenchyme, respectively (Fig. 2F). In stage-matched *ta^2^* MNPs, however, *SHH* expression was reduced and *PTCH1* was diffuse within the mesenchyme (Fig. 2G). Concordantly, *BMP4* and the downstream transcriptional target *MUSCLE SEGMENT HOMEOBOX 1* (*MSX1*; Barlow and Francis-West, 1997) were expanded into the oral mandibular epithelium and mesenchyme, respectively when compared to stage-matched controls (Fig. 2H-I). Additionally, as there was not a significant change in ectodermal *FGF8* expression, mesenchymal expression of its downstream transcriptional target *LIM HOMEOBOX 6* (*LHX6*; Grigoriou et al., 1998) was not significantly altered (Fig. 2J-K).

Following the observation that *BMP4* and its downstream target *MSX1* were upregulated in the early *ta^2^* MNP, we next sought to examine BMP pathway activity immediately prior to the onset of chondrogenesis. We performed scRNA-seq on HH26 control and *ta^2^* MNPs and utilized this dataset to run CellChat analysis to quantitatively analyze signal transduction pathways and infer cellular communication networks (Jin et al., 2021). In the control MNP, one epithelial cluster (cluster 1, outlined in blue in Fig. S4A) expressed *BMP4*, signaled to itself, as well as numerous neural crest mesenchyme and epithelial clusters (Fig. 3A-B). In the *ta^2^* MNP, however, a second, ectopic epithelial cell population (cluster 15, outlined in red in Fig. S4A) was identified in addition to that which was found in the control MNP (Fig. 3A-B). Cluster 15 both received a BMP signal from cluster 1 and signaled to the same epithelial and mesenchymal cell populations as cluster 1 (Fig. 3B). Interestingly, many neural crest mesenchyme and epithelial cell populations in the *ta^2^* mandibular prominence had an increased role for permitting (or acting as an “influencer” of) BMP signaling pathway activity (Fig. 3A). Furthermore, applying CellChat analytics to the Hedgehog and FGF signaling pathways revealed no significant changes between control and *ta^2^* MNPs as only a singular epithelial cluster (cluster 3, outlined in purple in Fig. S4A) sent and received a Hedgehog signal (Fig. S4B-C) and the FGF pathway was not detectable. Collectively, these results suggested the presence of ectopic cartilaginous processes in the *ta^2^* MNP correlated with ectopic BMP signaling activity at the level of the *BMP4* ligand expression.

**Figure 3.**
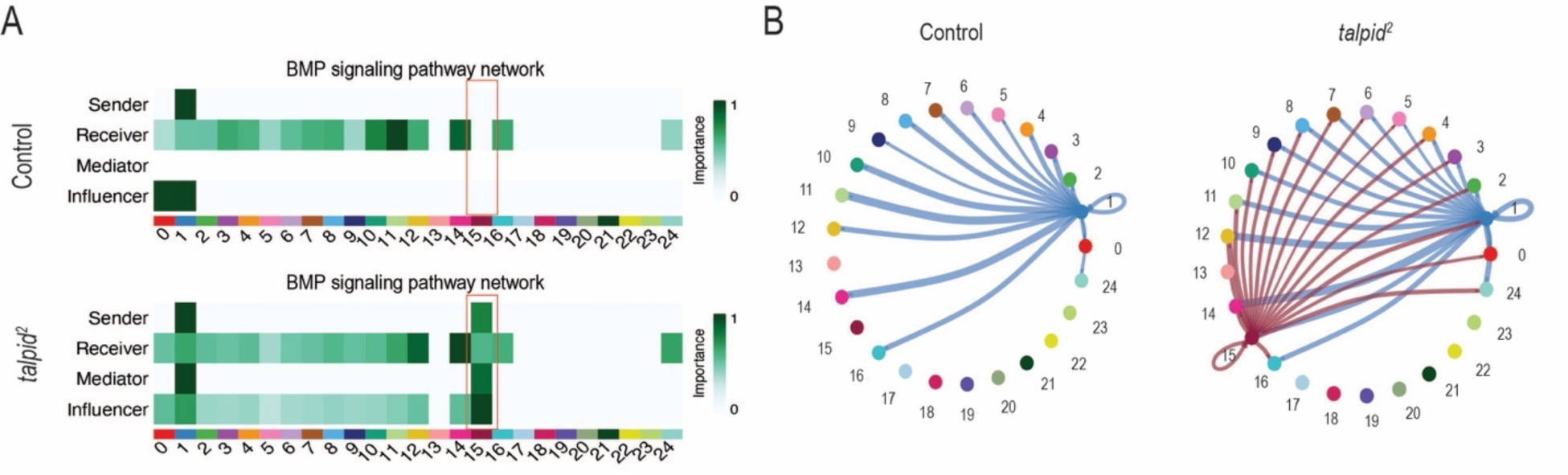
Cell-cell communication analysis reveals an ectopic BMP signaling center in the *ta^2^* MNP. (A-B) BMP signaling dynamics in HH26 control and *ta^2^* mandibular prominences as visualized via heatmap (A) and circle plot (B). Red boxes in panel A denote the ectopic BMP signaling center found in the single-cell transcriptomic dataset from the *ta^2^* mandibular prominence. The darker green shading in the heat map represents the greater role of a certain cell population. “Mediator” refers to the ability of a cell population to gatekeep information flow within a signaling pathway while “influencer” refers to the ability of a cell population to regulate a signaling pathway’s information flow.

### Conditional deletion of C2cd3 in murine models histologically and molecularly recapitulated ta^2^ cartilaginous phenotypes

Although our expression data and CellChat analysis suggested a role for epithelial signaling in the emergence of ectopic cartilage processes, the avian model system was not easily amenable to definitively testing this hypothesis. Since the *ta^2^ C2CD3* mutation was hypomorphic and affected all tissues (Chang et al., 2014), we transitioned to a murine model system and utilized three distinct Cre drivers to conditionally ablate *C2cd3* in tissues that contribute to the developing mandible. First, the *Wnt1-Cre2* driver recombined within CNCCs that comprise the craniofacial mesenchyme and differentiate into musculoskeletal tissues without the accompanying ectopic *Wnt1* expression of the widely-used *Wnt1-Cre* (Lewis et al., 2013). Second, the *Crect* driver recombined in the surface and oral ectoderm (Reid et al., 2011). Third, the *AP2-Cre* driver recombined in the surface and oral ectoderm, the CNCC-derived mesenchyme, and the neuroectoderm (Macatee et al., 2003). To determine the tissue-specific requirement for *C2cd3* function, we utilized previously generated mice with a *C2cd3^ex4-5flox^* allele (Chang et al., 2021) in combination with these Cre drivers (Fig. 4A-D) and assayed the resulting Meckel’s cartilage phenotype.

**Figure 4.**
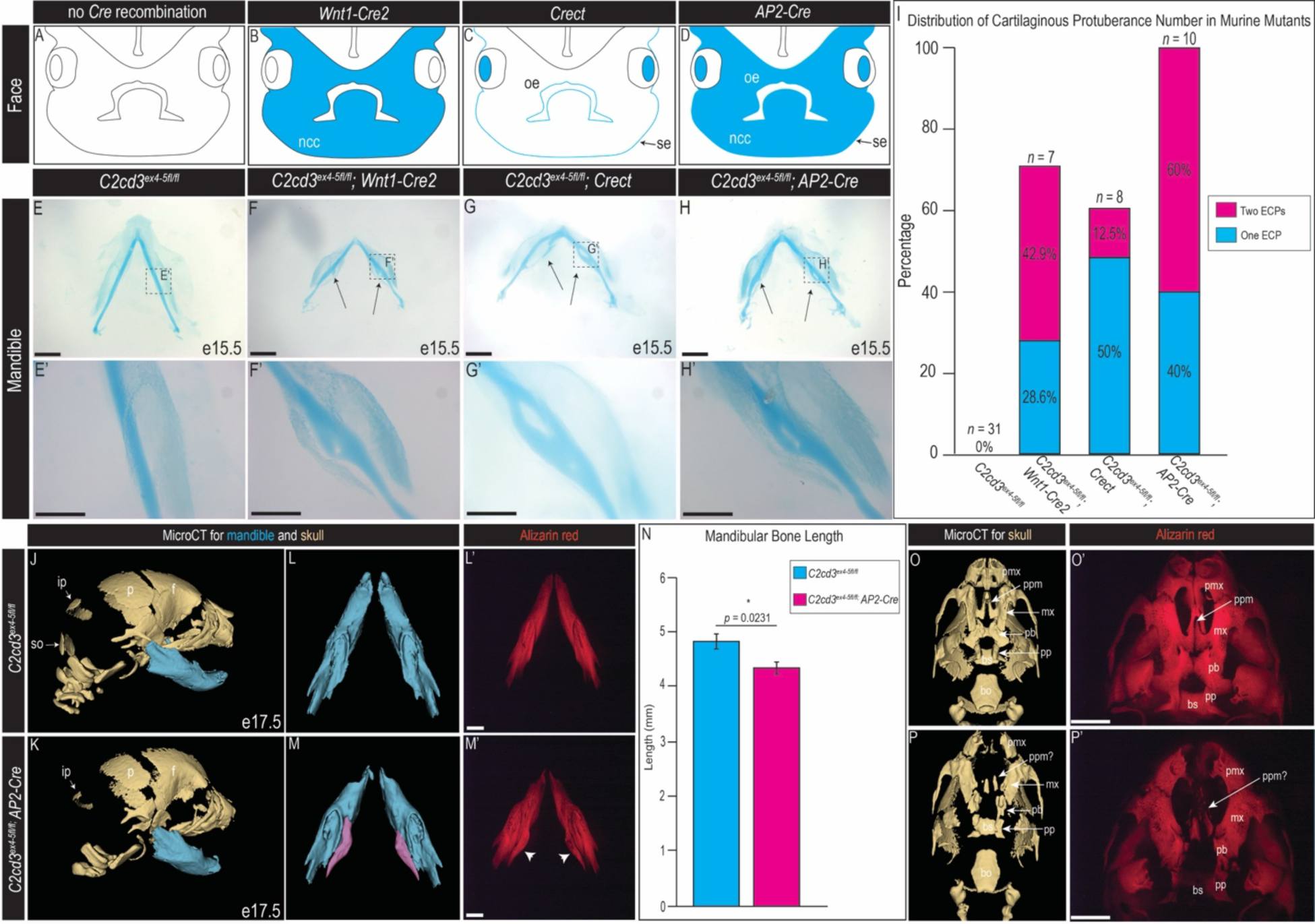
Conditional loss of *C2cd3* in murine epithelium and mesenchyme recapitulates *ta^2^* craniofacial phenotypes. (A-D) Schematization of spatial domains of Cre recombination (blue) for control (A), *Wnt1-Cre2* (B), *Crect* (C), and *AP2-Cre* (D) in e11.5 murine embryo faces. (E-H’) Wholemount Alcian blue staining for cartilage of e15.5 *C2cd3^ex4-5f/f^* (E, E’), *C2cd3^ex4-5f/f^; Wnt1-Cre2* (F, F’), *C2cd3^ex4-5f/f^; Crect* (G, G’), and *C2cd3^ex4-5f/f^; AP2-Cre* (H, H’) mandibles. Arrows in F, G, and H denote the presence of ectopic cartilaginous protuberances with images of higher magnification in F’, G’, and H’. (I) Bar chart of distribution of number of ectopic cartilaginous protuberances observed in *C2cd3* murine mutants. (J-K) Sagittal views of microCT-scanned E17.5 *C2cd3^ex4-5fl/fl^* (J, *n* = 3) and *C2cd3^ex4-5fl/fl^; AP2-Cre* (K, *n* = 3) heads with the mandible highlighted in blue. (L-M’) Dorsal views of E17.5 *C2cd3^ex4-5fl/fl^* (L-L’) and *C2cd3^ex4-5fl/fl^; AP2-Cre* (M-M’) mandibles from microCT scans (L, M) and Alizarin red staining (L’, M’). Purple shading in panel M and white arrowheads in M’ denote ectopic skeletal elements. (N) Morphometric analysis of mandibular bone lengths from microCT-scanned *C2cd3^ex4-5fl/fl^* and *C2cd3^ex4-5fl/fl^; AP2-Cre* mandibles. (O-P’) Ventral views of E17.5 *C2cd3^ex4-5fl/fl^* (O-O’) and *C2cd3^ex4-5fl/fl^; AP2-Cre* (P-P’) palates from microCT scans (O, P) and Alizarin red staining (O’, P’). bo: basioccipital, bs: basisphenoid, f: frontal bone, ip: interparietal bone, mx: maxilla, ncc: neural crest cells, oe: oral epithelium, p: parietal bone, pb: palatine bone, pmx: premaxilla, pp: pterygoid process, ppm: palatine process of the maxilla, se: surface ectoderm, so: supraoccipital. Asterisk denotes *p* < 0.05. Scale bars: 1mm (E, F, G, H, L’, M’, O’, P’), 500μm (E’, F’, G’, H’).

Alcian blue staining of E15.5 *C2cd3^ex4-5fl/fl^; Wnt1-Cre2* mandibles revealed that 48.9% (*n* = 3/7) of embryos presented with bilateral, ectopic processes on Meckel’s cartilage that resembled those present in the *ta^2^* (Fig. 4E-F’, I). Furthermore, 28.6% (*n* = 2/7) embryos presented with one, unilateral ectopic process and 28.6% (n = 2/7) embryos did not present with any ectopic process (Fig. 4I, Fig. S5A-B). *C2cd3^ex4-5fl/fl^; Crect* embryos presented with bilateral and unilateral ectopic processes in 12.5% (*n* = 1/8) and 50% (*n* = 4/8) of embryos, respectively (Fig. 4G-G’, I, Fig. S5C). The remaining *C2cd3^ex4-5fl/fl^; Crect* embryos (37.5%, *n* = 3/8) did not present with ectopic cartilaginous processes (Fig. S5D). Interestingly, *C2cd3^ex4-5fl/fl^; AP2-Cre* embryos presented with ectopic cartilaginous processes, either bilateral (60%, *n* = 6/10) or unilateral (40%, *n* = 4/10; Fig. 4H-I, Fig. S5E), at the highest level of penetrance of all Cre drivers. Thus, these data suggested an essential role for *C2cd3,* and cilia, in both epithelial and CNCC-derived mesenchymal tissues. Of note, loss of *C2cd3* in the *Scx-*lineage did not result in ectopic cartilage (Fig. S6), further suggesting that the onset of ectopic cartilage was a result of early aberrant mandibular patterning rather than altered fate decisions.

With the observation that conditional deletion of *C2cd3* in CNCC mesenchyme and craniofacial ectoderm in murine embryos phenocopied the *ta^2^* ectopic cartilage phenotype, we further characterized *C2cd3^ex4-5fl/fl^; AP2-Cre* embryos to understand if this model recapitulated *ta^2^* cranial phenotypes. microCT scans of *C2cd3^ex4-5fl/fl^; AP2-Cre* embryos revealed micrognathia as well as an ectopic mandibular skeletal element similar to that previously reported in *ta^2^* embryos (Fig. 4J-N) (Bonatto Paese et al., 2021a). In addition to the lower jaw phenotypes, *ta^2^* embryos also present with palatal clefting and hypoplastic palatine bones (Bonatto Paese et al., 2022; Chang et al., 2014; Schock et al., 2015). Indeed, *C2cd3^ex4-5fl/fl^; AP2-Cre* embryos presented with palatal clefting characterized by hypoplastic maxillary, premaxillary, and palatine bones as well as an absent palatine process and a dysmorphic pterygoid process (Fig. 4O-P’). These results further validated the importance of *C2cd3* and cilia in tissues of the craniofacial complex and suggested a conserved role between species.

With the ectopic cartilaginous processes in *C2cd3^ex4-5fl/fl^; AP2-Cre* resembling those in *ta^2^* embryos, we performed wholemount HCR to determine if the molecular basis for these phenotypes were conserved between species. As expected in e10.5 control MNPs, *Shh* was expressed in the oropharyngeal ectoderm, *Fgf8* was expressed in the proximal aboral ectoderm, and *Bmp4* was expressed in the distal aboral ectoderm (Fig. 5A). Similar to the *ta^2^*, *Bmp4* expression was expanded into the oral aspect, whereas *Shh* was diminished, and *Fgf8* was unchanged in e10.5 *C2cd3^ex4-5fl/fl^; AP2-Cre* MNPs (Fig. 5B). Sagittal views and quantification of expression domains of both *C2cd3^ex4-5fl/fl^* and *C2cd3^ex4-5fl/fl^; AP2-Cre* confirmed expansion of *Bmp4* into the oral ectoderm and reduction of *Shh* expression, with no significant change to *Fgf8* expression (Fig. 5C-E). Associated target gene expression was commensurately altered in CNCCs of *C2cd3^ex4-5fl/fl^; AP2-Cre* MNPs (Fig. 5F-K). These results ultimately suggested a conserved mechanism for the induction of ectopic cartilaginous processes between avian and murine embryos.

**Figure 5.**
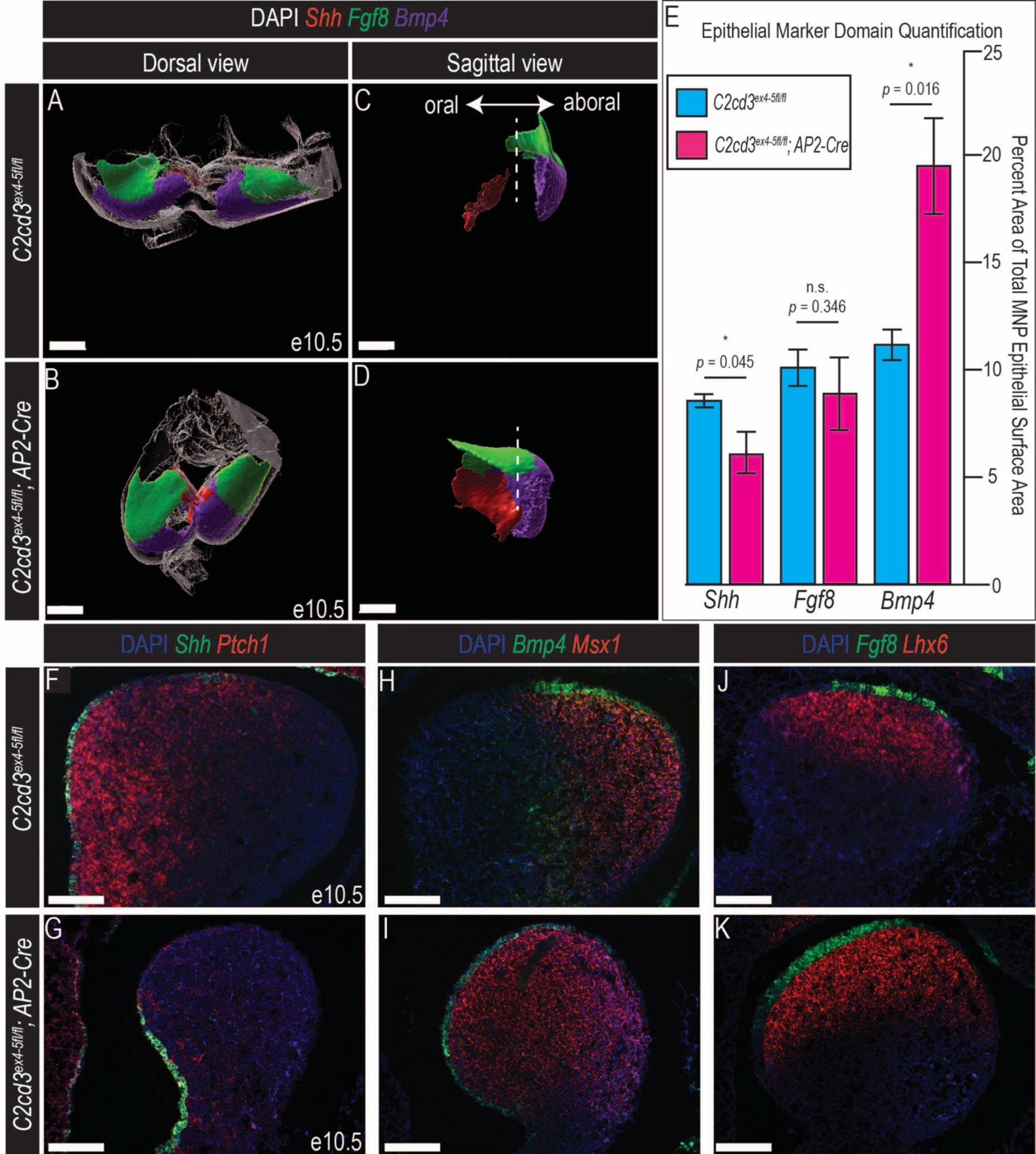
Conditional loss of C2cd3 in mandibular epithelium and mesenchyme recapitulates *ta^2^* molecular mandibular patterning. (A-D’) Three-dimensional renderings of wholemount HCR expression data for *Shh* (red), *Fgf8* (green), and *Bmp4* (purple) in e10.5 *C2cd3^ex4-5fl/fl^* (A, C, *n* = 3) and *C2cd3^ex4-5fl/fl^*; *AP2-Cre* (B, D, *n* = 3) MNPs. (E) Quantitation of three-dimensional surface renderings of *Shh*, *Fgf8*, and *Bmp4* in e10.5 *C2cd3^ex4-5fl/fl^* and *C2cd3^ex4-5fl/fl^*; *AP2-Cre* MNPs as a percentage of total mandibular surface area. (F-K) Section RNAscope *in situ* hybridization for *Shh* and *Ptch1* (F-G), *Bmp4* and *Msx1* (H-I), and *Fgf8* and *Lhx6* (J-K) on sagittal sections of e10.5 *C2cd3^ex4-5fl/fl^* (F, H, J, *n* = 3) and *C2cd3^ex4-5fl/fl^*; *AP2-Cre* (G, I, K, *n* = 3) MNPs. The dotted lines in C and D denote the midpoint of the oral-aboral axis of the MNP. Scale bars: 200μm (A-D), 100μm (F-K). *p* > 0.05, n.s. signifies no significance.

## Discussion

Mandibular dysmorphologies are a common feature of craniofacial ciliopathies. However, the mechanisms that govern mandibular musculoskeletal tissue patterning remains nebulous. Herein, we utilized both avian and murine model systems with mutations in *C2cd3* to understand how disruptions in ciliary function impacted signaling events that guide CNCC differentiation into musculoskeletal tissues. We found that the loss of *C2cd3*, and subsequent loss of ciliary extension, led to an ectopic epithelial BMP signaling center that propagated a BMP signal in the mandibular CNCC mesenchyme (Fig. 6). This study further expands our understanding of the necessity of primary cilia in regulating the spatial organization of signals that govern mandibular patterning.

**Figure 6.**
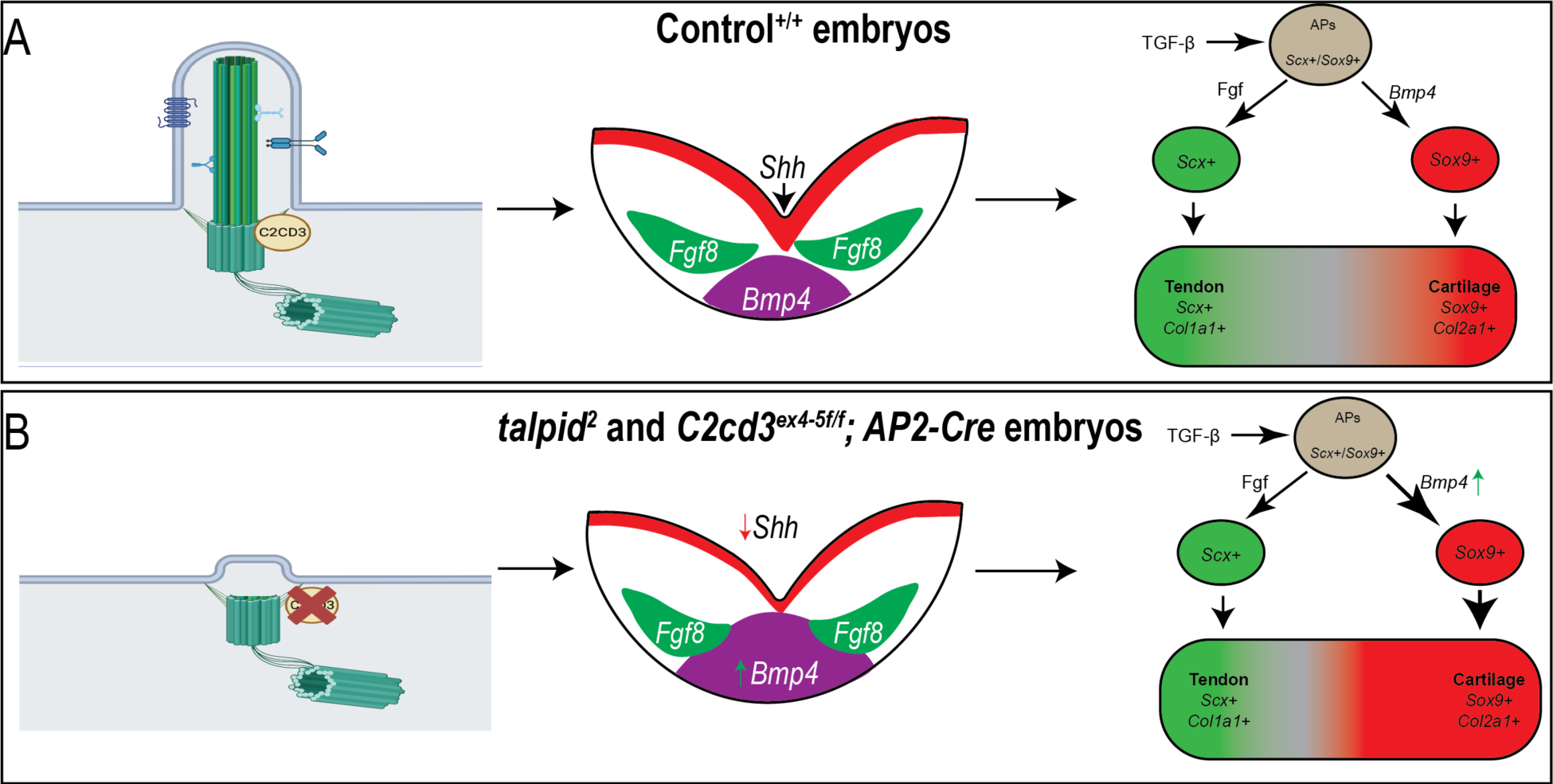
Summary of Meckel’s cartilage development in control and C2cd3 avian and murine mutant embryos. (A) In control conditions with normal C2cd3 expression and ciliogenesis, *Bmp4, Shh,* and Fgf8 expression is normal in the developing MNP. Normal patterning of the MNP results in normal differentiation of tendon and cartilage. (B) When C2cd3 is mutated, such as in *ta^2^* avian and *C2cd3^ex4-5fl/fl^; AP2-Cre* murine mutants, ciliogenesis is perturbed. There is a subsequent decrease in *Shh* expression and an increase in *Bmp4* expression in the developing mandibular prominence. Augmented *Bmp4* expression correlates with ectopic chondrogenesis during mandibular musculoskeletal differentiation. Cilium diagrams were created using BioRender.com.

### The primary cilium as a signaling hub for Hedgehog and BMP pathways

The cilium has long been associated with transduction of the Hh pathway due to localization of several pathway members to the cilium itself and the similarities of ciliopathic and Hedgehog related phenotypes (Goetz and Anderson, 2010; Huangfu et al., 2003; Huangfu and Anderson, 2005). Furthermore, the observed trafficking of Gli transcription factors through the cilium, which is essential for proper Hh signal transduction, has also been definitively and repeatedly documented (Haycraft et al., 2005; Kim et al., 2009; Wen et al., 2010). Interestingly, some but not all, of the mandibular (ectopic bone and expanded cartilage) and molecular (decreased Hh activity) readouts observed in both *ta^2^* and murine *C2cd3* mutant embryos were previously reported in conditional knockouts of the intraflagellar protein Ift88, and the Hh pathway transducer, Smoothened (Kitamura et al., 2020; Xu et al., 2019). Our RNAscope gene expression analysis supports the suggestion that reduced canonical Hh signaling may indeed contribute to the ectopic cartilages observed in our mutants; however, CellChat analysis did not validate that as the primary causal mechanism. Based on these data, we suggest that the unique phenotype may be associated with either a mutation in a protein localized to the centriole versus one localized to the axoneme, or additional aberrant signaling.

While the evidence for the relationship between the cilium and Hh has been observed and published by multiple groups, the evidence for if and how other molecular pathways (Wnt, FGF, BMP, PDGF, etc.) utilize the cilium has been less clear. In the case of BMP, current data suggests that TGFβ/BMP receptors are recruited to the cilium in order to activate SMADs (Anvarian et al., 2019). In addition to the localization of receptors to the ciliary membrane, proteins necessary for receptor recycling, pathway phosphorylation, and pathway feedback have all been reported to localize to the ciliary base/subdistal appendages (Clement et al., 2013; Miyazawa and Miyazono, 2017; Westlake et al., 2011). Since both RNAscope and CellChat analyses support increased BMP expression and signaling we hypothesize that the combinatorial impact of reduced Hh and increased BMP is causal for the ectopic cartilaginous processes, as well as accompanying musculoskeletal malformations observed in *ta^2^* and murine *C2cd3* mutant embryos. However, the exact mechanism of this combinatorial molecular insult is still unclear. It is possible that ectopic BMP expression and subsequent increased activity is secondary to reduced Hh activity since there is a well-documented antagonistic relationship between the two pathways (Bastida et al., 2009; Patten and Placzek, 2002; Xu et al., 2019). It is also possible that both pathways are directly impacted by a C2cd3 mutation. While we cannot eliminate the possibility of a BMP ligand being transduced through the cilium through a currently unknown mechanism, our molecular expression studies, in conjunction with other work utilizing ciliopathic animal models, favor a mechanism by which the cilium fine-tunes the antagonistic relationship between the BMP and Hh pathways during tissue patterning (Horner and Caspary, 2011; Kitamura et al., 2020). Future work into understanding if and how downstream factors of the Hh pathway mediate this antagonistic relationship, and potentially interact with phospho-Smad transcription factors, could provide this mechanistic insight.

### Is ectopic cartilage in talpid^2^ associated with atavism?

Previous studies characterizing *ta^2^* embryos revealed integumentary outgrowths on both the lower and upper jaws that resemble first-generation teeth from crocodilians as a result of shifts in the oral-aboral patterning molecules *SHH* and *BMP4* (Harris et al., 2006). The location of the ectopic cartilaginous processes in the distal portion of the mandible coincides with the emergence of archosaurian-like teeth in the *ta^2^*. Interestingly, secondary cartilages have been shown to emerge in sites of tooth eruption in the fossils of non-avian dinosaur hatchlings (Bailleul et al., 2013, 2012). While we are limited in our analysis of the *ta^2^* due to embryonic lethality, the altered musculoskeletal anatomy in the distal mandible at the ectopic cartilaginous processes does not exclude the possibility that they are associated with ectopic tooth formation.

Taking the presence of archosaurian-like teeth, together with the more recently discovered skeletal patterning in the limb that resembled ancestral tetrapods (Bonatto Paese et al., 2021b), our finding of ectopic cartilaginous processes that resemble entheses solidifies the *ta^2^* as an experimental model to dissect the molecular mechanisms underlying the emergence of atavisms, or recurrent ancestral characteristics, during skeletal development and evolution in comparison to embryos from reptilian and other avian species. Additional phenotyping efforts and examination of accompanying molecular changes in the *ta^2^* may reveal critical insights into the evolution of jaw, limb, and other musculoskeletal structures.

### Biomedical relevance of ectopic cartilage growth in ta^2^ for human ciliopathies

Craniofacial anomalies, including those present in ciliopathies, are among the most common congenital conditions affecting humans. Furthermore, they represent a significant biomedical burden of approximately $700 million per year. Surgical repair of craniofacial conditions is difficult and often requires a large source of skeletal tissue to reconstruct the facial skeleton. Depending upon the skeletal deficit, distinct treatment options are utilized. Common treatments for craniofacial skeletal anomalies include bone grafting with growth factor (typically BMP) supplementation and distraction osteogenesis. Bone grafting involves removal of bone from one part of the body and using it to repair the anomaly/injury. This is the preferred treatment when the bone deficit is relatively small. Distraction osteogenesis is a surgical procedure which involves cutting and slowly separating bone, allowing the bone healing process to fill in the gap (Ilizarov, 1988; McCarthy et al., 1992). This eliminates the need for taking bone from elsewhere but does require healthy soft tissue. Both approaches, however, have variable outcomes and can cause damaging off-target effects (e.g., relapse, nerve damage, infection, device failure, etc.) (Holloway et al., 2014; Kahn, 2014). Thus, there is a clear need for an improved approach towards surgical repair of craniofacial anomalies as well as metrics to determine which strategy is best.

Some ciliopathies have been reported to present with ectopic skeletal structures (Bonatto Paese et al., 2021a; Bredrup et al., 2011; Cela et al., 2018; Kitamura et al., 2020; Tabler et al., 2013; Zhang et al., 2011). Of those, bony protrusions, or tori particularly on the palate (torus palatinus) (Bredrup et al., 2011) or mandible (torus mandibularis) have been observed. Despite the common occurrence of these structures, there has been a significant knowledge gap in the understanding of their origin. Recently one hypothesis put forward suggests mandibular tori originate embryonically from a bending of Meckel’s cartilage that undergoes endochondral ossification at the level of the mental foramen (Rodríguez-Vázquez et al., 2013). A separate study examining the success of mandibular advancement devices, like those used in distraction osteogenesis, suggested that the presence of torus mandibularis almost tripled the likelihood of a successful response to distraction osteogenesis (Diaz de Teran et al., 2022). These data prompted two questions: 1) are the ectopic cartilaginous processes we observed embryonic precursors of mandibular tori, and 2) would the presence of ectopic skeletogenic processes in ciliopathy patients with micrognathia make them good candidates for distraction osteogenesis? While the early embryonic lethality of the *ta^2^* prevents us from addressing the former question, the latter may be addressed by careful phenotyping and documentation of outcome measures in ciliopathy patients. Such steps should be taken in an effort to best move forward with appropriate treatment options for this growing class of diseases.

## Supplemental Information

Supplementary figures for this manuscript include 6 figures.

## Author Contributions

Conceptualization: SAB; Methodology: ECB, SJYH, CLBP; Validation: ECB; Formal analysis: ECB, SJYH, SAB; Investigation: ECB, SJYH, CLBP, AAL, MAP; Resources: SAB; Writing – original draft preparation: ECB, SAB; Writing – review and editing: ECB, SAB; Visualization: ECB, SJYH, SAB; Supervision: SAB; Project administration: SAB; Funding acquisition: ECB, SAB.

## Data Availability

The single-cell transcriptomic dataset used in this manuscript can be found on FaceBase.org under the accession code FB00001245.

## Acknowledgments

We thank the University of California, Davis Avian Facility, Kevin Bellido, Jackie Pisenti and Dr. Mary Delany for maintenance and husbandry of the *talpid^2^* colony. Technical assistance for confocal image acquisition and analysis was given by Dr. Matt Kofron, Dr. Marina George, and Sarah McLeod of the Cincinnati Children’s Bio-Imaging and Analysis Facility [RRID: SCR_022628]. We acknowledge Dr. Xiangning (Sharon) Wang and Dr. Lisa Lemen of the University of Cincinnati Preclinical Imaging Core for microCT acquisition. Assistance with single-cell RNA sequencing preparation and analysis was respectively given by the Cincinnati Children’s Single Cell Genomics Core [RRID: SCR_022653] and Konrad Thorner of the Cincinnati Children’s Center for Stem Cell and Organoid Medicine. We thank Drs. Licia Selleri and Trevor Williams for supplying *Crect* mice, Drs. Amy Merrill-Brugger and Ronen Schweitzer for supplying *Scx-Cre* mice, and Dr. Katherine Woronowicz for providing technical guidance on avian paralysis experiments. We also thank members of the Brugmann lab as well as members of E.C.B.’s dissertation committee for helpful comments and feedback.

## References

1. Abbott, U., Taylor, L.W., Abplanalp, H., 1960. Studies with talpid2, an embryonic lethal of the fowl. J. Hered. 51, 195–202. 10.1093/oxfordjournals.jhered.a106988

2. Abbott, U., Taylor, L.W., Abplanalp, H., 1959. A 2nd Talpid-like mutation in the fowl. Poult. Sci. 38, 1185.

3. Abramyan, J., Richman, J.M., 2018. Craniofacial development: discoveries made in the chicken embryo. Int. J. Dev. Biol. 62, 97–107. 10.1387/ijdb.170321ja

4. Abzhanov, A., Tabin, C.J., 2004. Shh and Fgf8 act synergistically to drive cartilage outgrowth during cranial development. Dev. Biol. 273, 134–148. 10.1016/j.ydbio.2004.05.028

5. Adel Al-Lami, H., Barrell, W.B., Liu, K.J., 2016. Micrognathia in mouse models of ciliopathies. Biochem. Soc. Trans. 44, 1753–1759. 10.1042/BST20160241

6. Anvarian, Z., Mykytyn, K., Mukhopadhyay, S., Pedersen, L.B., Christensen, S.T., 2019. Cellular signalling by primary cilia in development, organ function and disease. Nat. Rev. Nephrol. 15, 199–219. 10.1038/s41581-019-0116-9

7. Bailleul, A.M., Hall, B.K., Horner, J.R., 2013. Secondary Cartilage Revealed in a Non-Avian Dinosaur Embryo. PLoS ONE 8, e56937. 10.1371/journal.pone.0056937

8. Bailleul, A.M., Hall, B.K., Horner, J.R., 2012. First Evidence of Dinosaurian Secondary Cartilage in the Post-Hatching Skull of Hypacrosaurus stebingeri (Dinosauria, Ornithischia). PLOS ONE 7, e36112. 10.1371/journal.pone.0036112

9. Baker, K., Beales, P.L., 2009. Making sense of cilia in disease: The human ciliopathies. Am. J. Med. Genet. C Semin. Med. Genet. 151C, 281–295. 10.1002/ajmg.c.30231

10. Barlow, A.J., Francis-West, P.H., 1997. Ectopic application of recombinant BMP-2 and BMP-4 can change patterning of developing chick facial primordia. Development 124, 391–398. 10.1242/dev.124.2.391

11. Bastida, M.F., Sheth, R., Ros, M.A., 2009. A BMP-Shh negative-feedback loop restricts Shh expression during limb development. Development 136, 3779–3789. 10.1242/dev.036418

12. Bi, W., Deng, J.M., Zhang, Z., Behringer, R.R., de Crombrugghe, B., 1999. Sox9 is required for cartilage formation. Nat. Genet. 22, 85–89. 10.1038/8792

13. Billmyre, K.K., Klingensmith, J., 2015. Sonic hedgehog from pharyngeal arch 1 epithelium is necessary for early mandibular arch cell survival and later cartilage condensation differentiation. Dev. Dyn. 244, 564–576. 10.1002/dvdy.24256

14. Bitgood, M.J., McMahon, A.P., 1995. Hedgehog and Bmp genes are coexpressed at many diverse sites of cell-cell interaction in the mouse embryo. Dev. Biol. 172, 126–138. 10.1006/dbio.1995.0010

15. Blitz, E., Sharir, A., Akiyama, H., Zelzer, E., 2013. Tendon-bone attachment unit is formed modularly by a distinct pool of Scx-and Sox9-positive progenitors. Development 140, 2680–2690. 10.1242/dev.093906

16. Blitz, E., Viukov, S., Sharir, A., Shwartz, Y., Galloway, J.L., Pryce, B.A., Johnson, R.L., Tabin, C.J., Schweitzer, R., Zelzer, E., 2009. Bone Ridge Patterning during Musculoskeletal Assembly Is Mediated through SCX Regulation of Bmp4 at the Tendon-Skeleton Junction. Dev. Cell 17, 861–873. 10.1016/j.devcel.2009.10.010

17. Bobzin, L., Roberts, R.R., Chen, H.-J., Crump, J.G., Merrill, A.E., 2021. Development and maintenance of tendons and ligaments. Development 148, dev186916. 10.1242/dev.186916

18. Boczek, N.J., Hopp, K., Benoit, L., Kraft, D., Cousin, M.A., Blackburn, P.R., Madsen, C.D., Oliver, G.R., Nair, A.A., Na, J., Bianchi, D.W., Beek, G., Harris, P.C., Pichurin, P., Klee, E.W., 2018. Characterization of three ciliopathy pedigrees expands the phenotype associated with biallelic C2CD3 variants. Eur. J. Hum. Genet. 26, 1797–1809. 10.1038/s41431-018-0222-3

19. Bonatto Paese, C.L., Brooks, E.C., Aarnio-Peterson, M., Brugmann, S.A., 2021a. Ciliopathic micrognathia is caused by aberrant skeletal differentiation and remodeling. Development 148, dev194175. 10.1242/dev.194175

20. Bonatto Paese, C.L., Chang, C.-F., Kristeková, D., Brugmann, S.A., 2022. Pharmacological intervention of the FGF-PTH axis as a potential therapeutic for craniofacial ciliopathies. Dis. Model. Mech. 15, dmm049611. 10.1242/dmm.049611

21. Bonatto Paese, C.L., Hawkins, M.B., Brugmann, S.A., Harris, M.P., 2021b. Atavisms in the avian hindlimb and early developmental polarity of the limb. Dev. Dyn. 250, 1358–1367. 10.1002/dvdy.318

22. Bredrup, C., Saunier, S., Oud, M.M., Fiskerstrand, T., Hoischen, A., Brackman, D., Leh, S.M., Midtbø, M., Filhol, E., Bole-Feysot, C., Nitschké, P., Gilissen, C., Haugen, O.H., Sanders, J.-S.F., Stolte-Dijkstra, I., Mans, D.A., Steenbergen, E.J., Hamel, B.C.J., Matignon, M., Pfundt, R., Jeanpierre, C., Boman, H., Rødahl, E., Veltman, J.A., Knappskog, P.M., Knoers, N.V.A.M., Roepman, R., Arts, H.H., 2011. Ciliopathies with skeletal anomalies and renal insufficiency due to mutations in the IFT-A gene WDR19. Am. J. Hum. Genet. 89, 634– 643. 10.1016/j.ajhg.2011.10.001

23. Brooks, E.C., Bonatto Paese, C.L., Carroll, A.H., Struve, J.N., Nagy, N., Brugmann, S.A., 2021. Mutation in the Ciliary Protein C2CD3 Reveals Organ-Specific Mechanisms of Hedgehog Signal Transduction in Avian Embryos. J. Dev. Biol. 9, 12. 10.3390/jdb9020012

24. Brugmann, S.A., Allen, N.C., James, A.W., Mekonnen, Z., Madan, E., Helms, J.A., 2010. A primary cilia-dependent etiology for midline facial disorders. Hum. Mol. Genet. 19, 1577–1592. 10.1093/hmg/ddq030

25. Caruccio, N.C., Martinez-Lopez, A., Harris, M., Dvorak, L., Bitgood, J., Simandl, B.K., Fallon, J.F., 1999. Constitutive activation of sonic hedgehog signaling in the chicken mutant talpid(2): Shh-independent outgrowth and polarizing activity. Dev. Biol. 212, 137–149. 10.1006/dbio.1999.9321

26. Cela, P., Hampl, M., Shylo, N.A., Christopher, K.J., Kavkova, M., Landova, M., Zikmund, T., Weatherbee, S.D., Kaiser, J., Buchtova, M., 2018. Ciliopathy Protein Tmem107 Plays Multiple Roles in Craniofacial Development. J. Dent. Res. 97, 108–117. 10.1177/0022034517732538

27. Chang, C.-F., Brown, K.M., Yang, Y., Brugmann, S.A., 2021. Centriolar Protein C2cd3 Is Required for Craniofacial Development. Front. Cell Dev. Biol. 9, 647391. 10.3389/fcell.2021.647391

28. Chang, C.-F., Schock, E.N., O’Hare, E.A., Dodgson, J., Cheng, H.H., Muir, W.M., Edelmann, R.E., Delany, M.E., Brugmann, S.A., 2014. The cellular and molecular etiology of the craniofacial defects in the avian ciliopathic mutant talpid2. Development 141, 3003– 3012. 10.1242/dev.105924

29. Choi, H.M.T., Calvert, C.R., Husain, N., Huss, D., Barsi, J.C., Deverman, B.E., Hunter, R.C., Kato, M., Lee, S.M., Abelin, A.C.T., Rosenthal, A.Z., Akbari, O.S., Li, Y., Hay, B.A., Sternberg, P.W., Patterson, P.H., Davidson, E.H., Mazmanian, S.K., Prober, D.A., van de Rijn, M., Leadbetter, J.R., Newman, D.K., Readhead, C., Bronner, M.E., Wold, B., Lansford, R., Sauka-Spengler, T., Fraser, S.E., Pierce, N.A., 2016. Mapping a multiplexed zoo of mRNA expression. Development 143, 3632–3637. 10.1242/dev.140137

30. Choi, H.M.T., Schwarzkopf, M., Fornace, M.E., Acharya, A., Artavanis, G., Stegmaier, J., Cunha, A., Pierce, N.A., 2018. Third-generation in situ hybridization chain reaction: multiplexed, quantitative, sensitive, versatile, robust. Development 145, dev165753. 10.1242/dev.165753

31. Clement, C.A., Ajbro, K.D., Koefoed, K., Vestergaard, M.L., Veland, I.R., Henriques de Jesus, M.P.R., Pedersen, L.B., Benmerah, A., Andersen, C.Y., Larsen, L.A., Christensen, S.T., 2013. TGF-β signaling is associated with endocytosis at the pocket region of the primary cilium. Cell Rep. 3, 1806–1814. 10.1016/j.celrep.2013.05.020

32. Cortés, C.R., McInerney-Leo, A.M., Vogel, I., Rondón Galeano, M.C., Leo, P.J., Harris, J.E., Anderson, L.K., Keith, P.A., Brown, M.A., Ramsing, M., Duncan, E.L., Zankl, A., Wicking, C., 2016. Mutations in human C2CD3 cause skeletal dysplasia and provide new insights into phenotypic and cellular consequences of altered C2CD3 function. Sci. Rep. 6, 24083. 10.1038/srep24083

33. Couly, G., Grapin-Botton, A., Coltey, P., Ruhin, B., Le Douarin, N.M., 1998. Determination of the identity of the derivatives of the cephalic neural crest: incompatibility between Hox gene expression and lower jaw development. Development 125, 3445–3459. 10.1242/dev.125.17.3445

34. Cserjesi, P., Brown, D., Ligon, K.L., Lyons, G.E., Copeland, N.G., Gilbert, D.J., Jenkins, N.A., Olson, E.N., 1995. Scleraxis: a basic helix-loop-helix protein that prefigures skeletal formation during mouse embryogenesis. Development 121, 1099–1110. 10.1242/dev.121.4.1099

35. de Beer, G.R., Barrington, E.J.W., 1934. The segmentation and chondrification of the skull of the duck. Philos. Trans. R. Soc. Lond. Ser. B Contain. Pap. Biol. Character 223, 411–467. 10.1098/rstb.1934.0009

36. Diaz de Teran, T., Muñoz, P., de Carlos, F., Macias, E., Cabello, M., Cantalejo, O., Banfi, P., Nicolini, A., Solidoro, P., Gonzalez, M., 2022. Mandibular Torus as a New Index of Success for Mandibular Advancement Devices. Int. J. Environ. Res. Public. Health 19, 14154. 10.3390/ijerph192114154

37. Dvorak, L., Fallon, J.F., 1991. Talpid2 mutant chick limb has anteroposterior polarity and altered patterns of programmed cell death. Anat. Rec. 231, 251–260. 10.1002/ar.1092310213

38. Edom-Vovard, F., Schuler, B., Bonnin, M.-A., Teillet, M.-A., Duprez, D., 2002. Fgf4 positively regulates scleraxis and tenascin expression in chick limb tendons. Dev. Biol. 247, 351–366. 10.1006/dbio.2002.0707

39. Elliott, K.H., Brugmann, S.A., 2019. Sending mixed signals: Cilia-dependent signaling during development and disease. Dev. Biol. 447, 28–41. 10.1016/j.ydbio.2018.03.007

40. Elliott, K.H., Chen, X., Salomone, J., Chaturvedi, P., Schultz, P.A., Balchand, S.K., Servetas, J.D., Zuniga, A., Zeller, R., Gebelein, B., Weirauch, M.T., Peterson, K.A., Brugmann, S.A., 2020. Gli3 utilizes Hand2 to synergistically regulate tissue-specific transcriptional networks. eLife 9, e56450. 10.7554/eLife.56450

41. Fedorov, A., Beichel, R., Kalpathy-Cramer, J., Finet, J., Fillion-Robin, J.-C., Pujol, S., Bauer, C., Jennings, D., Fennessy, F., Sonka, M., Buatti, J., Aylward, S., Miller, J.V., Pieper, S., Kikinis, R., 2012. 3D Slicer as an Image Computing Platform for the Quantitative Imaging Network. Magn. Reson. Imaging 30, 1323–1341. 10.1016/j.mri.2012.05.001

42. Ferrante, M.I., Romio, L., Castro, S., Collins, J.E., Goulding, D.A., Stemple, D.L., Woolf, A.S., Wilson, S.W., 2009. Convergent extension movements and ciliary function are mediated by ofd1, a zebrafish orthologue of the human oral-facial-digital type 1 syndrome gene. Hum. Mol. Genet. 18, 289–303. 10.1093/hmg/ddn356

43. Franco, B., Thauvin-Robinet, C., 2016. Update on oral-facial-digital syndromes (OFDS). Cilia 5, 12. 10.1186/s13630-016-0034-4

44. Goetz, S.C., Anderson, K.V., 2010. The Primary Cilium: A Signaling Center During Vertebrate Development. Nat. Rev. Genet. 11, 331–344. 10.1038/nrg2774

45. Goodrich, L.V., Johnson, R.L., Milenkovic, L., McMahon, J.A., Scott, M.P., 1996. Conservation of the hedgehog/patched signaling pathway from flies to mice: induction of a mouse patched gene by Hedgehog. Genes Dev. 10, 301–312. 10.1101/gad.10.3.301

46. Grigoriou, M., Tucker, A.S., Sharpe, P.T., Pachnis, V., 1998. Expression and regulation of Lhx6 and Lhx7, a novel subfamily of LIM homeodomain encoding genes, suggests a role in mammalian head development. Development 125, 2063–2074. 10.1242/dev.125.11.2063

47. Hall, B.K., 2015. Bones and Cartilage: Developmental and Evolutionary Skeletal Biology, 2nd ed. Academic Press.

48. Hall, B.K., 1975. A simple, single-injection method for inducing long-term paralysis in embryonic chicks, and preliminary observations on growth of the tibia. Anat. Rec. 181, 767–777. 10.1002/ar.1091810408

49. Hamburger, V., Hamilton, H.L., 1951. A series of normal stages in the development of the chick embryo. J. Morphol. 88, 49–92. 10.1002/jmor.1050880104

50. Hao, Y., Hao, S., Andersen-Nissen, E., Mauck, W.M., Zheng, S., Butler, A., Lee, M.J., Wilk, A.J., Darby, C., Zager, M., Hoffman, P., Stoeckius, M., Papalexi, E., Mimitou, E.P., Jain, J., Srivastava, A., Stuart, T., Fleming, L.M., Yeung, B., Rogers, A.J., McElrath, J.M., Blish, C.A., Gottardo, R., Smibert, P., Satija, R., 2021. Integrated analysis of multimodal single-cell data. Cell 184, 3573–3587.e29. 10.1016/j.cell.2021.04.048

51. Harris, M.P., Hasso, S.M., Ferguson, M.W.J., Fallon, J.F., 2006. The Development of Archosaurian First-Generation Teeth in a Chicken Mutant. Curr. Biol. 16, 371–377. 10.1016/j.cub.2005.12.047

52. Havis, E., Bonnin, M.-A., Esteves de Lima, J., Charvet, B., Milet, C., Duprez, D., 2016. TGFβ and FGF promote tendon progenitor fate and act downstream of muscle contraction to regulate tendon differentiation during chick limb development. Development 143, 3839–3851. 10.1242/dev.136242

53. Haworth, K.E., Healy, C., Morgan, P., Sharpe, P.T., 2004. Regionalisation of early head ectoderm is regulated by endoderm and prepatterns the orofacial epithelium. Development 131, 4797–4806. 10.1242/dev.01337

54. Haworth, K.E., Wilson, J.M., Grevellec, A., Cobourne, M.T., Healy, C., Helms, J.A., Sharpe, P.T., Tucker, A.S., 2007. Sonic hedgehog in the pharyngeal endoderm controls arch pattern via regulation of Fgf8 in head ectoderm. Dev. Biol. 303, 244–258. 10.1016/j.ydbio.2006.11.009

55. Haycraft, C.J., Banizs, B., Aydin-Son, Y., Zhang, Q., Michaud, E.J., Yoder, B.K., 2005. Gli2 and Gli3 localize to cilia and require the intraflagellar transport protein polaris for processing and function. PLoS Genet. 1, e53. 10.1371/journal.pgen.0010053

56. Holloway, J.L., Ma, H., Rai, R., Burdick, J.A., 2014. Modulating hydrogel crosslink density and degradation to control bone morphogenetic protein delivery and in vivo bone formation. J. Control. Release Off. J. Control. Release Soc. 191, 63–70. 10.1016/j.jconrel.2014.05.053

57. Hoover, A.N., Wynkoop, A., Zeng, H., Jia, J., Niswander, L.A., Liu, A., 2008. C2cd3 is required for cilia formation and Hedgehog signaling in mouse. Development 135, 4049–4058. 10.1242/dev.029835

58. Horner, V.L., Caspary, T., 2011. Disrupted dorsal neural tube BMP signaling in the cilia mutant Arl13b hnn stems from abnormal Shh signaling. Dev. Biol. 355, 43–54. 10.1016/j.ydbio.2011.04.019

59. Hu, D., Colnot, C., Marcucio, R.S., 2008. The effect of BMP signaling on development of the jaw skeleton. Dev. Dyn. 237, 3727–3737. 10.1002/dvdy.21781

60. Hu, D., Marcucio, R.S., Helms, J.A., 2003. A zone of frontonasal ectoderm regulates patterning and growth in the face. Development 130, 1749–1758. 10.1242/dev.00397

61. Huangfu, D., Anderson, K.V., 2005. Cilia and Hedgehog responsiveness in the mouse. Proc. Natl. Acad. Sci. U. S. A. 102, 11325–11330. 10.1073/pnas.0505328102

62. Huangfu, D., Liu, A., Rakeman, A.S., Murcia, N.S., Niswander, L., Anderson, K.V., 2003. Hedgehog signalling in the mouse requires intraflagellar transport proteins. Nature 426, 83–87. 10.1038/nature02061

63. Ilizarov, G.A., 1988. The principles of the Ilizarov method. Bull. Hosp. Jt. Dis. Orthop. Inst. 48, 1– 11.

64. Jeong, J., Mao, J., Tenzen, T., Kottmann, A.H., McMahon, A.P., 2004. Hedgehog signaling in the neural crest cells regulates the patterning and growth of facial primordia. Genes Dev. 18, 937–951. 10.1101/gad.1190304

65. Jin, S., Guerrero-Juarez, C.F., Zhang, L., Chang, I., Ramos, R., Kuan, C.-H., Myung, P., Plikus, M.V., Nie, Q., 2021. Inference and analysis of cell-cell communication using CellChat. Nat. Commun. 12, 1088. 10.1038/s41467-021-21246-9

66. Kahn, M., 2014. Can we safely target the WNT pathway? Nat. Rev. Drug Discov. 13, 513–532. 10.1038/nrd4233

67. Kantaputra, P., Dejkhamron, P., Sittiwangkul, R., Katanyuwong, K., Ngamphiw, C., Sonsuwan, N., Intachai, W., Tongsima, S., Beales, P.L., Buranaphatthana, W., 2023. Dental Anomalies in Ciliopathies: Lessons from Patients with BBS2, BBS7, and EVC2 Mutations. Genes 14, 84. 10.3390/genes14010084

68. Kim, J., Kato, M., Beachy, P.A., 2009. Gli2 trafficking links Hedgehog-dependent activation of Smoothened in the primary cilium to transcriptional activation in the nucleus. Proc. Natl. Acad. Sci. U. S. A. 106, 21666–21671. 10.1073/pnas.0912180106

69. Kitamura, A., Kawasaki, M., Kawasaki, K., Meguro, F., Yamada, A., Nagai, T., Kodama, Y., Trakanant, S., Sharpe, P.T., Maeda, T., Takagi, R., Ohazama, A., 2020. Ift88 is involved in mandibular development. J. Anat. 236, 317–324. 10.1111/joa.13096

70. Klingenberg, C.P., Navarro, N., 2012. Development of the mouse mandible: a model system for complex morphological structures, in: Piálek, J., Macholán, M., Munclinger, P., Baird, S.J.E. (Eds.), Evolution of the House Mouse, Cambridge Studies in Morphology and Molecules: New Paradigms in Evolutionary Bio. Cambridge University Press, Cambridge, pp. 135–149. 10.1017/CBO9781139044547.008

71. Kolpakova-Hart, E., Jinnin, M., Hou, B., Fukai, N., Olsen, B.R., 2007. Kinesin-2 Controls Development and Patterning of the Vertebrate Skeleton by Hedgehog-and Gli3-Dependent Mechanisms. Dev. Biol. 309, 273–284. 10.1016/j.ydbio.2007.07.018

72. Köntges, G., Lumsden, A., 1996. Rhombencephalic neural crest segmentation is preserved throughout craniofacial ontogeny. Development 122, 3229–3242. 10.1242/dev.122.10.3229

73. Kosher, R.A., Kulyk, W.M., Gay, S.W., 1986. Collagen gene expression during limb cartilage differentiation. J. Cell Biol. 102, 1151–1156.

74. Lewis, A.E., Vasudevan, H.N., O’Neill, A.K., Soriano, P., Bush, J.O., 2013. The widely used Wnt1-Cre transgene causes developmental phenotypes by ectopic activation of Wnt signaling. Dev. Biol. 379, 229–234. 10.1016/j.ydbio.2013.04.026

75. Macatee, T.L., Hammond, B.P., Arenkiel, B.R., Francis, L., Frank, D.U., Moon, A.M., 2003. Ablation of specific expression domains reveals discrete functions of ectoderm-and endoderm-derived FGF8 during cardiovascular and pharyngeal development. Development 130, 6361–6374. 10.1242/dev.00850

76. Marigo, V., Scott, M.P., Johnson, R.L., Goodrich, L.V., Tabin, C.J., 1996. Conservation in hedgehog signaling: induction of a chicken patched homolog by Sonic hedgehog in the developing limb. Development 122, 1225–1233. 10.1242/dev.122.4.1225

77. McCarthy, J.G., Schreiber, J., Karp, N., Thorne, C.H., Grayson, B.H., 1992. Lengthening the human mandible by gradual distraction. Plast. Reconstr. Surg. 89, 1–8.

78. Miyazawa, K., Miyazono, K., 2017. Regulation of TGF-β Family Signaling by Inhibitory Smads. Cold Spring Harb. Perspect. Biol. 9, a022095. 10.1101/cshperspect.a022095

79. Muntifering, M., Castranova, D., Gibson, G.A., Meyer, E., Kofron, M., Watson, A.M., 2018. Clearing for Deep Tissue Imaging. Curr. Protoc. Cytom. 86, e38. 10.1002/cpcy.38

80. Murray, P.D.F., 1963. Adventitious (Secondary) cartilage in the chick, and the development of certain bones and articulation in the chick skull. Aust. J. Zool. 11, 368–430. 10.1071/zo9630368

81. Nagy, A., Gertsenstein, M., Vintersten, K., Behringer, R., 2009. Alcian blue staining of the mouse fetal cartilaginous skeleton. Cold Spring Harb. Protoc. 2009, pdb.prot5169. 10.1101/pdb.prot5169

82. Neubüser, A., Peters, H., Balling, R., Martin, G.R., 1997. Antagonistic Interactions between FGF and BMP Signaling Pathways: A Mechanism for Positioning the Sites of Tooth Formation. Cell 90, 247–255. 10.1016/S0092-8674(00)80333-5

83. Nonaka, K., Shum, L., Takahashi, I., Takahashi, K., Ikura, T., Dashner, R., Nuckolls, G.H., Slavkin, H.C., 1999. Convergence of the BMP and EGF signaling pathways on Smad1 in the regulation of chondrogenesis. Int. J. Dev. Biol. 43, 795–807.

84. Patten, I., Placzek, M., 2002. Opponent activities of Shh and BMP signaling during floor plate induction in vivo. Curr. Biol. CB 12, 47–52. 10.1016/s0960-9822(01)00631-5

85. Reid, B.S., Yang, H., Melvin, V.S., Taketo, M.M., Williams, T., 2011. Ecotdermal Wnt/β-Catenin signaling shapes the mouse face. Dev. Biol. 349, 261–269. 10.1016/j.ydbio.2010.11.012

86. Reiter, J.F., Leroux, M.R., 2017. Genes and molecular pathways underpinning ciliopathies. Nat. Rev. Mol. Cell Biol. 18, 533–547. 10.1038/nrm.2017.60

87. Roberts, R.R., Bobzin, L., Teng, C.S., Pal, D., Tuzon, C.T., Schweitzer, R., Merrill, A.E., 2019. FGF signaling patterns cell fate at the interface between tendon and bone. Development 146, dev170241. 10.1242/dev.170241

88. Robson, L.G., 1993. Cellular patterning of fast and slow fibres in the intermandibularis muscle of chick embryos. Development 117, 329–339. 10.1242/dev.117.1.329

89. Rodríguez-Vázquez, J.F., Sakiyama, K., Verdugo-López, S., Amano, O., Murakami, G., Abe, S., 2013. Origin of the torus mandibularis: an embryological hypothesis. Clin. Anat. N. Y. N 26, 944–952. 10.1002/ca.22275

90. Schindelin, J., Arganda-Carreras, I., Frise, E., Kaynig, V., Longair, M., Pietzsch, T., Preibisch, S., Rueden, C., Saalfeld, S., Schmid, B., Tinevez, J.-Y., White, D.J., Hartenstein, V., Eliceiri, K., Tomancak, P., Cardona, A., 2012. Fiji: an open-source platform for biological-image analysis. Nat. Methods 9, 676–682. 10.1038/nmeth.2019

91. Schneider, R.A., Hu, D., Helms, J.A., 1999. From head to toe: conservation of molecular signals regulating limb and craniofacial morphogenesis. Cell Tissue Res. 296, 103–109. 10.1007/s004410051271

92. Schock, E.N., Brugmann, S.A., 2017. Discovery, Diagnosis, and Etiology of Craniofacial Ciliopathies. Cold Spring Harb. Perspect. Biol. 9, a028258. 10.1101/cshperspect.a028258

93. Schock, E.N., Chang, C.-F., Struve, J.N., Chang, Y.-T., Chang, J., Delany, M.E., Brugmann, S.A., 2015. Using the avian mutant talpid2 as a disease model for understanding the oral-facial phenotypes of oral-facial-digital syndrome. Dis. Model. Mech. 8, 855–866. 10.1242/dmm.020222

94. Schweitzer, R., Chyung, J.H., Murtaugh, L.C., Brent, A.E., Rosen, V., Olson, E.N., Lassar, A., Tabin, C.J., 2001. Analysis of the tendon cell fate using Scleraxis, a specific marker for tendons and ligaments. Development 128, 3855–3866. 10.1242/dev.128.19.3855

95. Schweitzer, R., Zelzer, E., Volk, T., 2010. Connecting muscles to tendons: tendons and musculoskeletal development in flies and vertebrates. Development 137, 2807–2817. 10.1242/dev.047498

96. Semba, I., Nonaka, K., Takahashi, I., Takahashi, K., Dashner, R., Shum, L., Nuckolls, G.H., Slavkin, H.C., 2000. Positionally-dependent chondrogenesis induced by BMP4 is co-regulated by sox9 and msx2. Dev. Dyn. 217, 401–414. 10.1002/(SICI)1097-0177(200004)217:4<401::AID-DVDY7>3.0.CO;2-D

97. Shibata, S., Suda, N., Yoda, S., Fukuoka, H., Ohyama, K., Yamashita, Y., Komori, T., 2004. Runx2-deficient mice lack mandibular condylar cartilage and have deformed Meckel’s cartilage. Anat. Embryol. (Berl.) 208, 273–280. 10.1007/s00429-004-0393-2

98. Shigetani, Y., Nobusada, Y., Kuratani, S., 2000. Ectodermally Derived FGF8 Defines the Maxillomandibular Region in the Early Chick Embryo: Epithelial–Mesenchymal Interactions in the Specification of the Craniofacial Ectomesenchyme. Dev. Biol. 228, 73–85. 10.1006/dbio.2000.9932

99. Solem, R.C., Eames, B.F., Tokita, M., Schneider, R.A., 2011. Mesenchymal and mechanical mechanisms of secondary cartilage induction. Dev. Biol. 356, 28–39. 10.1016/j.ydbio.2011.05.003

100. Sugimoto, Y., Takimoto, A., Akiyama, H., Kist, R., Scherer, G., Nakamura, T., Hiraki, Y., Shukunami, C., 2013. Scx+/Sox9+ progenitors contribute to the establishment of the junction between cartilage and tendon/ligament. Development 140, 2280–2288. 10.1242/dev.096354

101. Sunadome, K., Erickson, A.G., Kah, D., Fabry, B., Adori, C., Kameneva, P., Faure, L., Kanatani, S., Kaucka, M., Dehnisch Ellström, I., Tesarova, M., Zikmund, T., Kaiser, J., Edwards, S., Maki, K., Adachi, T., Yamamoto, T., Fried, K., Adameyko, I., 2023. Directionality of developing skeletal muscles is set by mechanical forces. Nat. Commun. 14, 3060. 10.1038/s41467-023-38647-7

102. Svandova, E., Anthwal, N., Tucker, A.S., Matalova, E., 2020. Diverse Fate of an Enigmatic Structure: 200 Years of Meckel’s Cartilage. Front. Cell Dev. Biol. 8, 821. 10.3389/fcell.2020.00821

103. Tabler, J.M., Barrell, W.B., Szabo-Rogers, H.L., Healy, C., Yeung, Y., Perdiguero, E.G., Schulz, C., Yannakoudakis, B.Z., Mesbahi, A., Wlodarczyk, B., Geissmann, F., Finnell, R.H., Wallingford, J.B., Liu, K.J., 2013. Fuz Mutant Mice Reveal Shared Mechanisms between Ciliopathies and FGF-Related Syndromes. Dev. Cell 25, 623–635. 10.1016/j.devcel.2013.05.021

104. Thauvin-Robinet, C., Lee, J.S., Lopez, E., Herranz-Pérez, V., Shida, T., Franco, B., Jego, L., Ye, F., Pasquier, L., Loget, P., Gigot, N., Aral, B., Lopes, C.A.M., St-Onge, J., Bruel, A.-L., Thevenon, J., González-Granero, S., Alby, C., Munnich, A., Vekemans, M., Huet, F., Fry, A.M., Saunier, S., Rivière, J.-B., Attié-Bitach, T., Garcia-Verdugo, J.M., Faivre, L., Mégarbané, A., Nachury, M.V., 2014. The oral-facial-digital syndrome gene C2CD3 encodes a positive regulator of centriole elongation. Nat. Genet. 46, 905–911. 10.1038/ng.3031

105. Tucker, A.S., Matthews, K.L., Sharpe, P.T., 1998. Transformation of tooth type induced by inhibition of BMP signaling. Science 282, 1136–1138. 10.1126/science.282.5391.1136

106. Tucker, A.S., Yamada, G., Grigoriou, M., Pachnis, V., Sharpe, P.T., 1999. Fgf-8 determines rostral-caudal polarity in the first branchial arch. Development 126, 51–61. 10.1242/dev.126.1.51

107. Vainio, S., Karavanova, I., Jowett, A., Thesleff, I., 1993. Identification of BMP-4 as a signal mediating secondary induction between epithelial and mesenchymal tissues during early tooth development. Cell 75, 45–58. 10.1016/S0092-8674(05)80083-2

108. Wang, F., Flanagan, J., Su, N., Wang, L.-C., Bui, S., Nielson, A., Wu, X., Vo, H.-T., Ma, X.-J., Luo, Y., 2012. RNAscope: A novel in situ RNA analysis platform for formalin-fixed, paraffin-embedded tissues. J. Mol. Diagn. 14, 22–29. 10.1016/j.jmoldx.2011.08.002

109. Wen, X., Lai, C.K., Evangelista, M., Hongo, J.-A., de Sauvage, F.J., Scales, S.J., 2010. Kinetics of Hedgehog-Dependent Full-Length Gli3 Accumulation in Primary Cilia and Subsequent Degradation. Mol. Cell. Biol. 30, 1910–1922. 10.1128/MCB.01089-09

110. Westlake, C.J., Baye, L.M., Nachury, M.V., Wright, K.J., Ervin, K.E., Phu, L., Chalouni, C., Beck, J.S., Kirkpatrick, D.S., Slusarski, D.C., Sheffield, V.C., Scheller, R.H., Jackson, P.K., 2011. Primary cilia membrane assembly is initiated by Rab11 and transport protein particle II (TRAPPII) complex-dependent trafficking of Rabin8 to the centrosome. Proc. Natl. Acad. Sci. U. S. A. 108, 2759–2764. 10.1073/pnas.1018823108

111. Woronowicz, K.C., Gline, S.E., Herfat, S.T., Fields, A.J., Schneider, R.A., 2018. FGF and TGFβ signaling link form and function during jaw development and evolution. Dev. Biol. 444, S219–S236. 10.1016/j.ydbio.2018.05.002

112. Wright, W.E., Sassoon, D.A., Lin, V.K., 1989. Myogenin, a factor regulating myogenesis, has a domain homologous to MyoD. Cell 56, 607–617. 10.1016/0092-8674(89)90583-7

113. Xu, J., Liu, H., Lan, Y., Adam, M., Clouthier, D.E., Potter, S., Jiang, R., 2019. Hedgehog signaling patterns the oral-aboral axis of the mandibular arch. eLife 8, e40315. 10.7554/eLife.40315

114. Ye, X., Zeng, H., Ning, G., Reiter, J.F., Liu, A., 2014. C2cd3 is critical for centriolar distal appendage assembly and ciliary vesicle docking in mammals. Proc. Natl. Acad. Sci. U. S. A. 111, 2164–2169. 10.1073/pnas.1318737111

115. Zhang, Z., Wlodarczyk, B.J., Niederreither, K., Venugopalan, S., Florez, S., Finnell, R.H., Amendt, B.A., 2011. Fuz Regulates Craniofacial Development through Tissue Specific Responses to Signaling Factors. PLoS ONE 6, e24608. 10.1371/journal.pone.0024608

