## Supplementary figures and images for "The ciliary protein C2cd3 is required for mandibular musculoskeletal tissue patterning"

### Figure S1

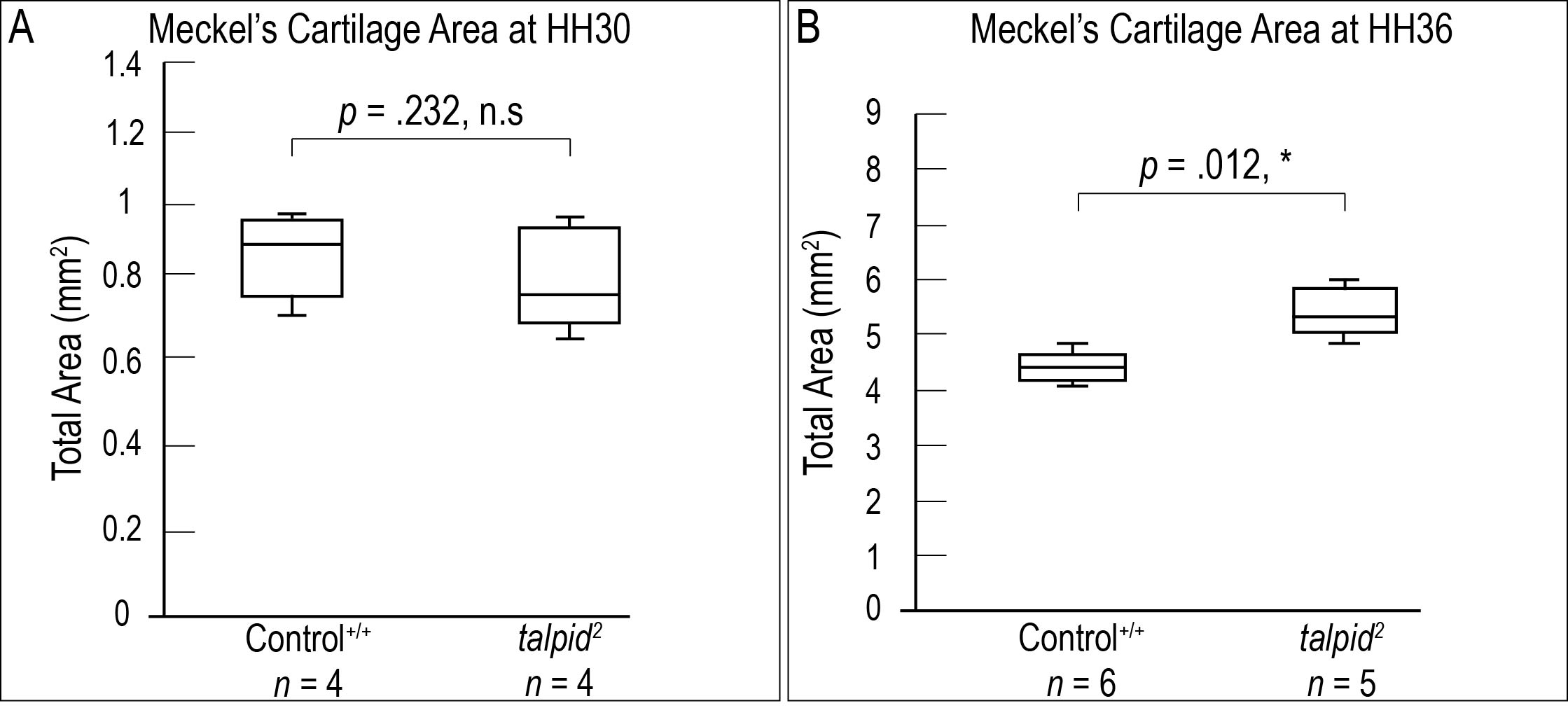

### Figure S2

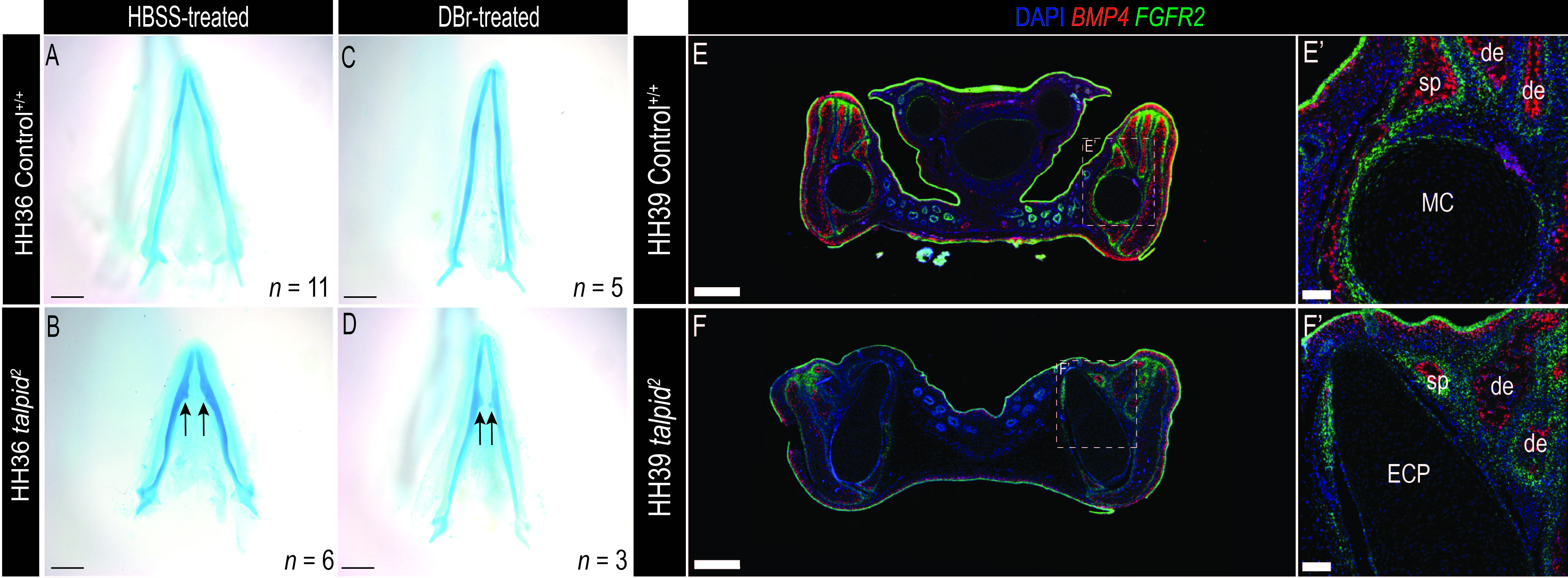

### Figure S3

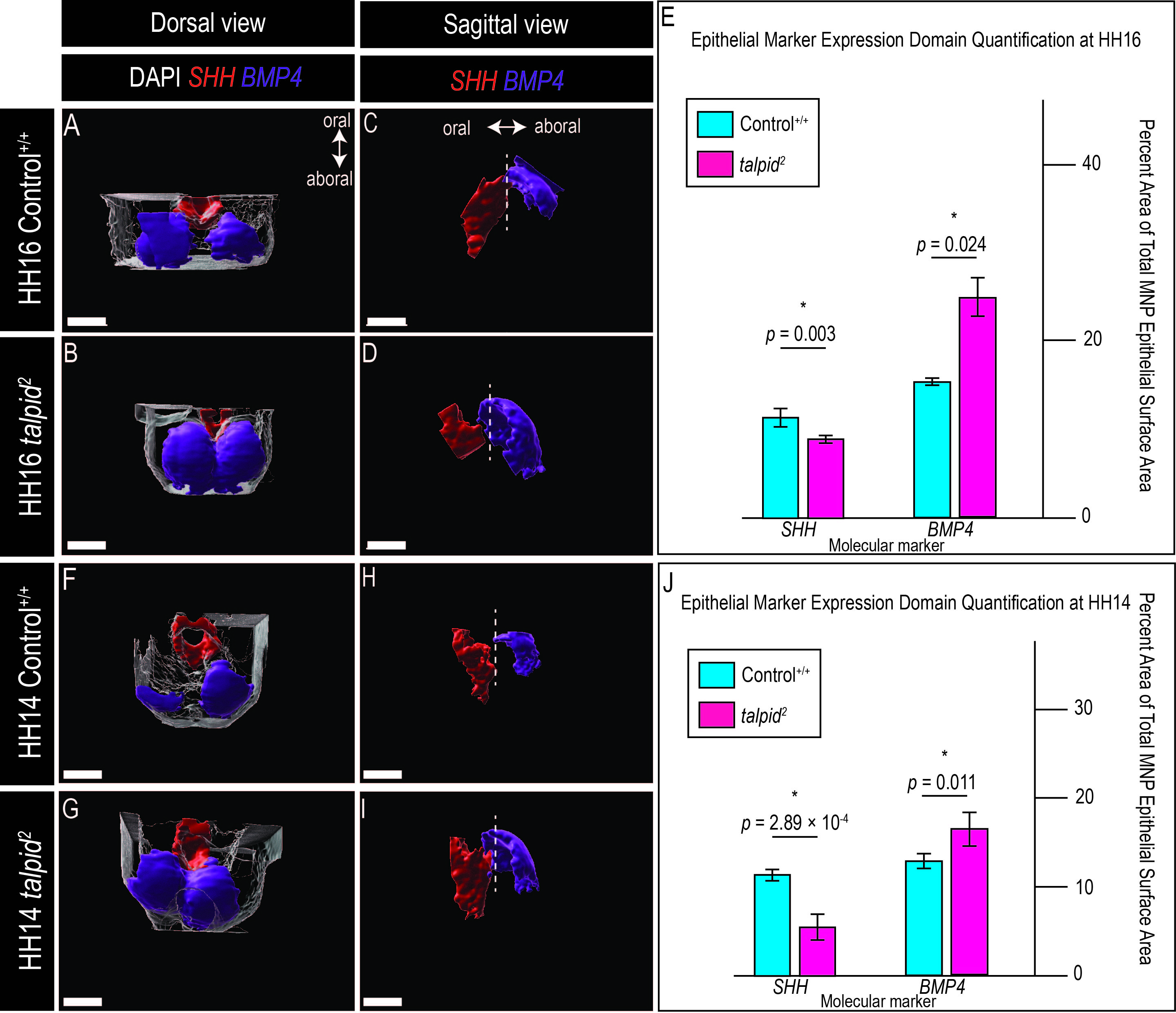

### Figure S4

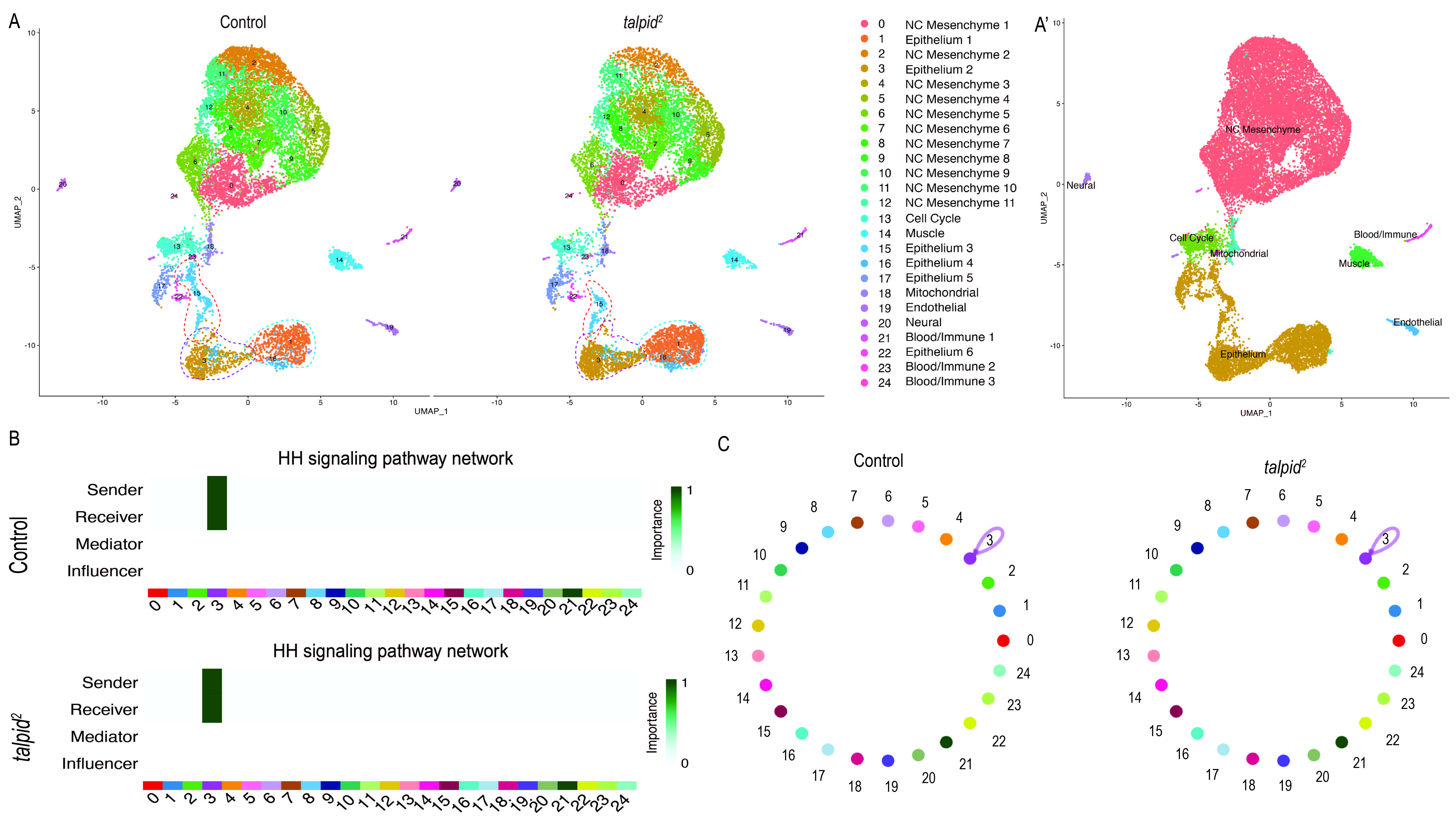

### Figure S5

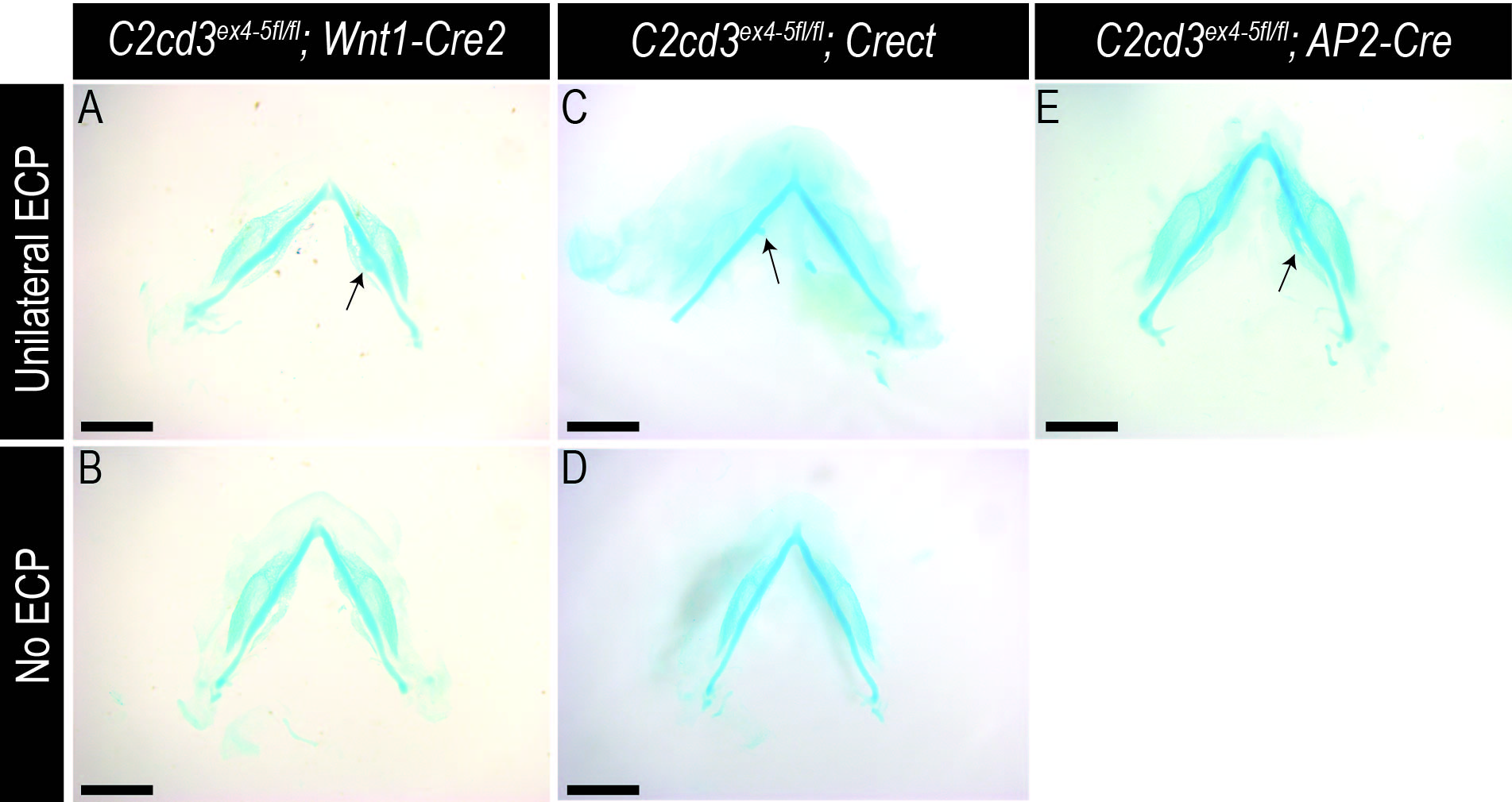

### Figure S6

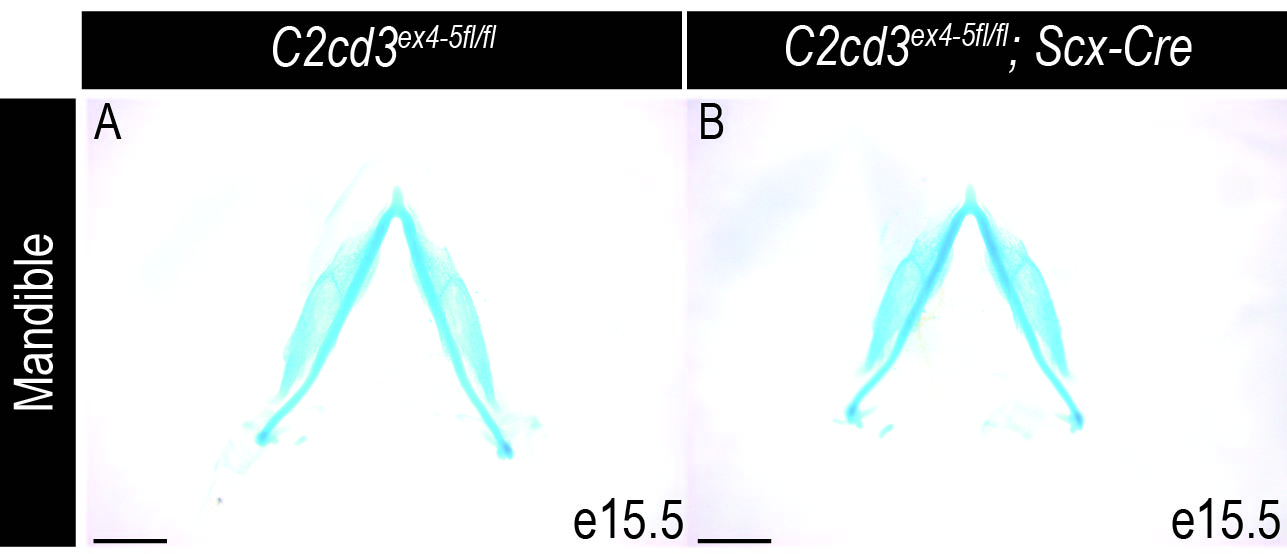
